# NMR Spectroscopy for the Validation of AlphaFold2 Structures

**DOI:** 10.1101/2025.02.04.636507

**Authors:** Jake Williams, Isabelle A. Gagnon, Joseph R. Sachleben

## Abstract

The introduction of AlphaFold has fundamentally changed our ability to predict the structure of proteins from their primary sequence of amino acids. As machine learning (ML) and artificial intelligence (AI) based protein prediction continues to advance, we examine the potential of hybrid techniques that combine experiment and computation that may yield more accurate structures than AI alone with significantly reduced experimental burden. We have developed heuristics comparing N-edited NOESY spectra and AlphaFold predicted structures that seek to determine whether the predicted structure reasonably describes the structure of the protein which generated the NOESY. These heuristics are fundamentally different than those used before in that they concentrate on interresidue contacts and not specific HH pairs. We present a large collection of data connecting entries across the BMRB, PDB and AlphaFold Database that includes experimentally derived structures and corresponding spectra, establishing it as a means to develop and test hybrid methods utilizing AlphaFold and NMR spectra to perform structure determination. These data test the new heuristics’ ability to identify inaccurate AlphaFold structures. A support vector machine was developed to test the consistency of NMR data with predicted structure and we show its application to the structure of an unsolved engineered protein, LoTOP.

## 1 Introduction

The rise of big data and deep learning methods have fundamentally changed the way in which we pursue science. For example, it has long been a goal in biochemistry to predict the 3D structure, or fold, of a protein from its primary amino acid sequence. [1, 2]. AlphaFold [3], a deep learning model for exactly this, has revolutionized our ability to make such predictions. This was first demonstrated in a community experiment called the Critical Assessment of Structure Prediction (CASP), which compares computational methods for determining protein structures from their amino acid sequences. AlphaFold2 competed in CASP-14 where it was found to be ‘competitive with experimental accuracy’ [4].

While AlphaFold2 is a powerful system, the protein folding problem is not completely solved, nor has deep learning been shown to be entirely self-sufficient. For example, AlphaFold2 is known to fail on certain point mutations [5] and unusual sequences [6, 7], cannot model proteins that have multiple stable structures [8], does not model the dynamics of systems in solution or reactions [9], and struggles with complexes and multimers [10]. AlphaFold2 does work to mitigate its own potential pitfalls by augmenting its predictions with a model of the uncertainty involved in those predictions [3]. In many of these areas, new systems have been developed from the AlphaFold framework. Recently, AlphaFold-Multimer [11] and AlphaFold3 [12] were introduced to tackle the challenges involved in predicting protein complexes. The success and predictive power of AlphaFold2 have led to a range of competitors. For example, RoseTTAFold is a competing neural network based method that has shown similar success [13]. We expect that the methods presented here using AlphaFold2 will be relevant to all future prediction methods.

AlphaFold models have been compared to NMR structures in a series of papers by Montelione and coworkers. [14, 15] They have shown in a small data set of nine protein targets, that the AlphaFold2 predicted structures fit remarkably well with NMR data. Güntert and coworkers showed that with accurate AlphaFold models, NMR peak assignment can be particularly efficient and accurate.[16] Neither of these groups looked at how to use NMR data to determine if the predicted structure is consistent with NMR data, which is the focus of this paper.

Understanding the structures predicted by AlphaFold, including when they are accurate or inaccurate, has become a focal point for those studying protein structure determination. We seek to generate hybrid methods that combine traditional experimental methods and current deep learning methods to balance the trade offs between the convenience of deep learning models and time consuming and expensive experimental structural determination. We propose to combine nuclear magnetic resonance (NMR) experiments with deep-learning models to check and refine the modeled protein structures. This paper concentrates on using Nuclear Overhauser Effect Spectroscopy (NOESY) to validate structures through the pairwise atomic distance restraints it imposes [17]. Even in settings with reduced information about the nature of the peaks generated in a NOESY spectrum, the experimental data can be used to determine likelihood of structure quality [18], and these methods inspire our work. Other NMR experiments which probe characteristics such as rigidity [19], bond torsion angles[20],and residual dipolar couplings can also be used to validate structures and will be the subject of future work.

This paper focuses on experimentally validating predicted protein structures with constraints derived from easily acquired NMR spectra; i.e., the ^15^N edited NOESY spectrum. In order to quantify the agreement between predicted structure and NMR data, two heuristics have been developed; the Contact and Distance Scores, CS and DS, respectively. These heuristics are based on the idea of ‘potential contacts’, as described fully in section 2.2. These heuristics are tested on a few example structures to demonstrate their feasibility. We have mined data from multiple sources to compile a dataset that contains proteins with experimentally derived structures, experimental NMR parameters, and AlphaFold2 predicted structures. This data collection is an important tool for creating and testing hybrid methods and forms the basis of this and future work. It allows us to test our heuristics over our current state of knowledge. We introduce a predictive model, a support vector machine (SVM), called Structural Prediction Assessment by NMR (SPANR), that was trained on our dataset and tests the accuracy of the AlphaFold predicted structures relative to the experimentally derived structures. We demonstrate SPANR’s ability to distinguish high-quality predictions and use SPANR to test the AlphaFold structure of a novel engineered protein, LoTOP.

## 2 Computational Methods

### 2.1 Collecting Existing Data

Publicly available archived data were collected and used in this analysis. There were three primary sources of these data: the Biological Magnetic Resonance Data Bank (BMRB), the AlphaFold DB, and Protein Data Bank (PDB). These sources provided NMR assignments, predicted protein structures, and NMR solved protein structures, respectively, and are the primary inputs to our data pipeline, shown as the top row in figure 1. We explicitly eliminate membrane proteins, where high-quality NMR data are difficult to acquire, and proteins measured in 8M urea solution.

**Fig. 1:**
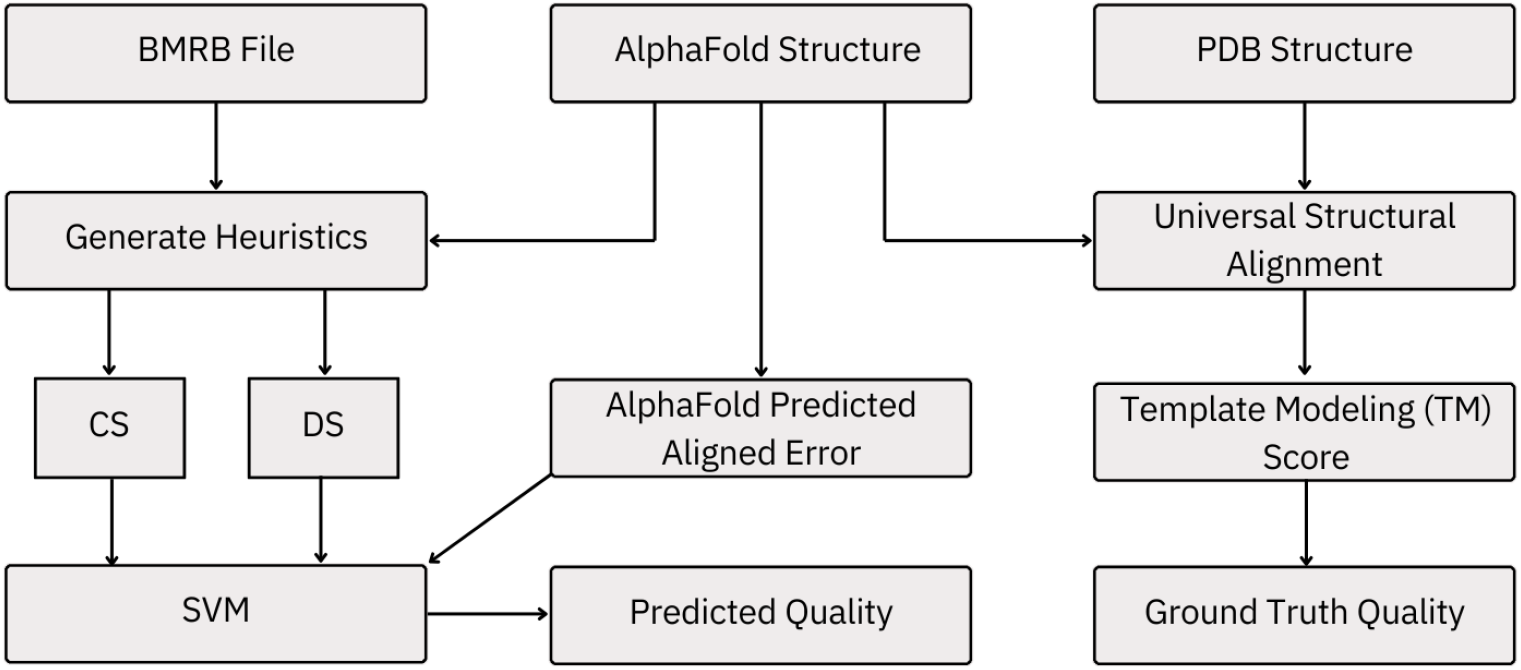
Visualization of the flow of data and analysis. Our model compares spectra from the BMRB to structures predicted by AlphaFold using the heuristics described in section 2.2. These heuristics and the predicted uncertainty from AlphaFold feed a small support vector machine which predicts the quality of the AlphaFold structure. The true quality of the AlphaFold structure is determined by the TM-score between the AlphaFold structure and the experimentally derived structure in the PDB.

#### 2.1.1 BMRB

The Biological Magnetic Resonance Data Bank (BMRB) supplied our experimental NMR data [21]. The BMRB is a repository for spectroscopic results obtained in magnetic resonance experiments, and contains experimental NMR spectra, lists of assigned chemical shift values for individual atoms, as well as some lists of peaks directly from the spectra. Data are archived in STAR format, which is a dictionary-based method for data exchange commonly used in molecular structure sciences [22, 23].

These files were screened for lists of assigned chemical shifts and NOESY peaks; however, the majority of data reported do not contain the NOESY data. Excluding files without NOESY data would produces a dataset with only 117 members, which is too small to produce reliable results, but does allow us to test our results. For this reason, we opted to simulate all NOESY spectra using the assigned chemical shifts and the PDB structure. This process is described in section 2.1.3.

#### 2.1.2 PDB

The NMR data acquired from the BMRB are associated with experimentally derived structures in the Protein Data Bank (PDB), the primary repository for all 3D structures of proteins, nucleic acids, and complexes [24]. AlphaFold also obtained its training data from the PDB.

We have focused primarily on protein structures that were solved by NMR spectroscopy. Thus, if the BMRB file lists only PDB structures that were solved using crystallography or another non-NMR method, we discard this structure from our dataset. Even though AlphaFold was primarily trained on protein structures derived from X-ray diffraction data, we decided to compare its ability to find the structure in solution as determined by NMR spectroscopy where artifacts due to effects such as crystal packing are absent. PDB files provide the atomic coordinates of each atom in an XYZ format. When the positions of hydrogen atoms were excluded from the PDB files, RDKit was used to calculate their positions [25]. We assume that the NMR determined PDB structure is the ground truth for each protein in our dataset.

#### 2.1.3 NOESY Simulation

From the list of assigned chemical shifts and an experimentally determined structure, we simulate the peaks from an ^15^N-edited NOESY experiment, Algorithm 1. The algorithm loops over all backbone amide hydrogens as the ‘base’ hydrogens and calculates the distance to all other hydrogens using the first model of the NMR-determined structure in the PDB file. No attempts are made to include the effects of motion on the NOEs. We add a peak to the spectrum with a probability that is related to the inverse of the distance to the 6th power, as shown in line 12. The inverse 6th power is related to the intensity of the NOE.[26]

This simulation algorithm allows us to approximate the list of peaks we might see in a 3D ^15^N-edited NOESY spectrum. However, it does not require the actual list of NOESY peaks or any parsing of the BMRB file beyond the assigned chemical shifts, which are the most reliably available and easily parsed section of the BMRB files. We are able to do these simulations because we have identified a ground truth 3D structure from the PDB. We used this simulation as a final check of the validity of the experimental data, removing any structures for which less than half of the residues have generated a potential contact, as these data are not informative enough to make conclusions about the full structure.

Importantly, this method of simulating the NOESY spectrum does not produce an idealized, “perfect” spectrum. Due to the probability distribution used, many NOESY peaks are missing and a few unreasonable peaks are added. If two hydrogen atoms are in steric contact with an interresidue distance of 2.4Å, a peak will always be included in our simulated spectrum; however, if the HH distance increases to 4Å, the peak will only be included 50% of the time. There is also a 2% chance that an unreasonable NOESY peak with an 8Å internuclear distance will be included. Thus, these simulated data are not idealized but are missing many potentially observable NOESY peaks and contain some spurious peaks making the simulated data a reasonable proxy for experimentally acquired data. In the supplemental information, we used the 117 members of our dataset that contained experimental NOESY data to demonstrate that the simulation algorithm accurately recapitulates experimental NOESY lists. We found that on average 91% of the simulated peaks were also in the experimental data, 9% of the simulated peaks were added and not seen in the experimental data, and 36% of the experimental peaks were missing from the simulated list. This demonstrates that our simulated data were far from idealized data and provide a rigorous test of our heuristics and appropriate training data for SPANR. Histograms of these comparisons are in the supplemental information.

#### 2.1.4 AlphaFold DB

The predicted structures were collected from the AlphaFold Database, [27], which contains over 200 million predicted structures. Each entry in the database consists of a single predicted conformer for its amino acid sequence. Each record in the database also contains the “predicted aligned error” (PAE), which is a matrix that describes the uncertainty in each pairwise distance between residues.[3]

We correlate the AlphaFold DB entries with the BMRB files using the UniProt accession code, which is the protein sequence identifier used by AlphaFold DB. For each structure, we obtain the AlphaFold2 best predicted structure and the PAE. We verify that the amino acid sequence in the AlphaFold DB matches the amino acid sequence in the BMRB file. Using the AlphaFold DB to collect predicted structures allows us to examine larger amounts of data than would otherwise be possible.

#### 2.1.5 Data Collection Summary

Our final dataset has 3085 structures, each with an experimentally derived ground-truth PDB structure, an AlphaFold predicted structure, and assigned NMR chemical shifts from which we simulate a list of NOESY peaks. In the rest of this paper we will refer to this dataset as the “original” dataset, as it has not been augmented with other data.

Among the 3085 members of our data set, 117 have experimental NOESY data associated with their NMR determined PDB structures. These data were used to test our simulated NOESY data and the results that use simulated NOESY spectra.

### 2.2 Analysis Techniques

We have developed a set of heuristics that take as input a structure and a list of NOESY peaks and output a single value intended to quantify the likelihood that the structure could have generated the NOESY peaks. The input structure can derive from any source: structurally determined by X-ray crystallography, NMR, or cryo-EM, or predicted by AlphaFold or any other tool. The heuristics can be applied to almost any type of protein for which experimental NMR data exist, and even unstructured regions of proteins can give useful insight for distinguishing high-quality from lower-quality structures by providing negative examples.

The quality of the predicted structure is quantified by calculating the TM-score[28] between it and the PDB structure. We judge our heuristics to be useful if they correlate well with the TM-score. Most powerfully in terms of efficiency, these heuristics are calculated without explicitly assigning the NOESY peaks. This is done using a set of ‘potential contacts’, which is described in section 2.2.1. Potential contacts feed into the two heuristics, CS and DS, described in sections 2.2.2 and 2.2.3, respectively.

#### 2.2.1 Potential Contacts

Potential contacts are derived from the ^15^N-edited NOESY spectrum and the ^1^H assignments. The ^15^N-edited NOESY spectrum provides ^1^H chemical shifts of hydrogens that are near the hydrogen of the assigned amide, HN. Due to the extensive overlap of the ^1^H chemical shifts and the limited resolution of the spectra, only potential contacts between HN and the other hydrogen atoms can be determined. A potential contact is determined if the chemical shift difference between the NOESY peak and its possible assignment is less than Δ_*shift*_ ppm. Thus, each off-diagonal NOESY peak provides a list of potential contacts to the residue assigned to the given HN peak, Figure 2. This uncertainty in assignment leads to ambiguous NOEs in our analysis. It should be emphasized that our potential contacts are between residues and not specific HH pairs. The specific H atom on the residue is not used in our analysis, only the residue identity.

**Fig. 2:**
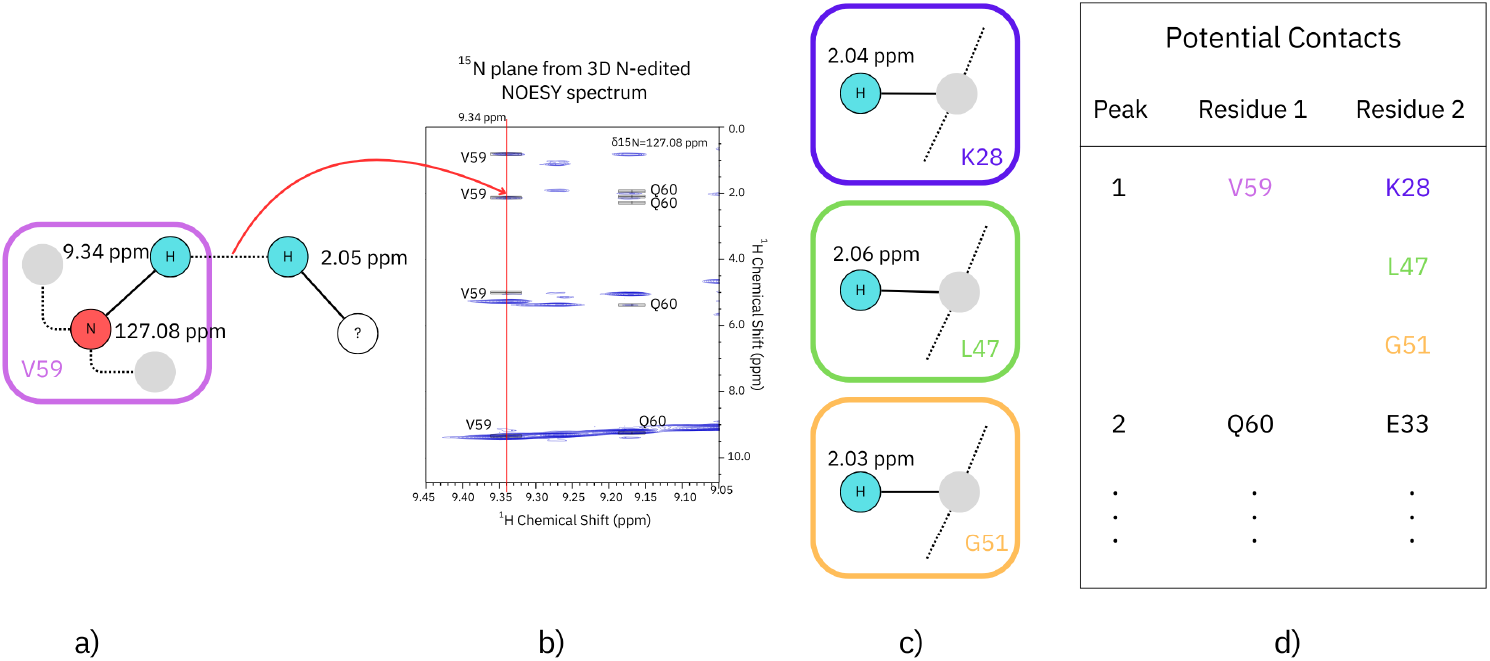
Potential Contacts are generated by comparing the chemical shifts of the assigned H’s with the positions of the off-diagonal peaks in the NOESY spectrum. a) shows the chemical shifts of the HN and N of V59 of a protein. b) shows an *δ*_*N*_ = 127.08 ppm plane of an ^15^N-edited NOESY spectrum. ^1^H peaks along the red line at 9.34 ppm are either the HN peak of V59 or due to H’s near to it. One such peak is at 2.05 ppm. c) Several H’s have shifts near to 2.05 and are potential contacts to V59.

Ambiguous assignments have been used in previous validation methods. In particular, Huang et al. descibe using recall, precision, and F-measure (RPF) in NMR structure validation verifying that the proposed structure is consistent with the observed NOESY peaks.[18] We have similarly used RPF to compare our NOESY and predicted structure interresidue contacts. These results shown in the SI as well as the resulting SVMs trained with these addition heuristics. In the next two sections, we will describe the algorithms for our own methods of using these potential contacts to assess the quality of a predicted structure.

#### 2.2.2 Contact Score

CS compares the interresidue contacts in the predicted structure to those predicted by the NMR data. It concentrates only on long-range contacts; i.e., those whose primary sequence separation is greater than 4 residues. As described above, NOESY peaks in the N-edited NOESY spectrum provide potential contacts which are compared to those seen the predicted structure. Scoring these potential contacts both in terms of the number of contacts and their agreement with their assignment determines CS.

This procedure is illustrated in Figure 3A and detailed in Algorithm 2. The spatial distance between C_*α*_ of residues identified by the potential contacts is determined from the proposed structure and compared to an upper cutoff threshold *ϵ*_*CS*_. *ϵ*_*CS*_ was treated as a hyperparameter and optimized by a grid search, which determined that for two residues to be “in contact” their C_*α*_s had to be within 12Å. The grid search optimized the correlation between CS and the TM-score, with full details in the SI. If the spatial distance between the two CA atoms in residues i and j is less than *ϵ*_*CS*_ and these residues are separated by at least 5 amino acids, are deemed to be “in contact” and a score is calculated. This score measures the interresidue contact weighted by the chemical shift agreement. It is calculated from the chemical shift difference, *δ*_*P C*_, and the chemical shift tolerance, *δ*_*measure*_, such that a score between 0 and 1 is assigned for every contact.

**Fig. 3:**
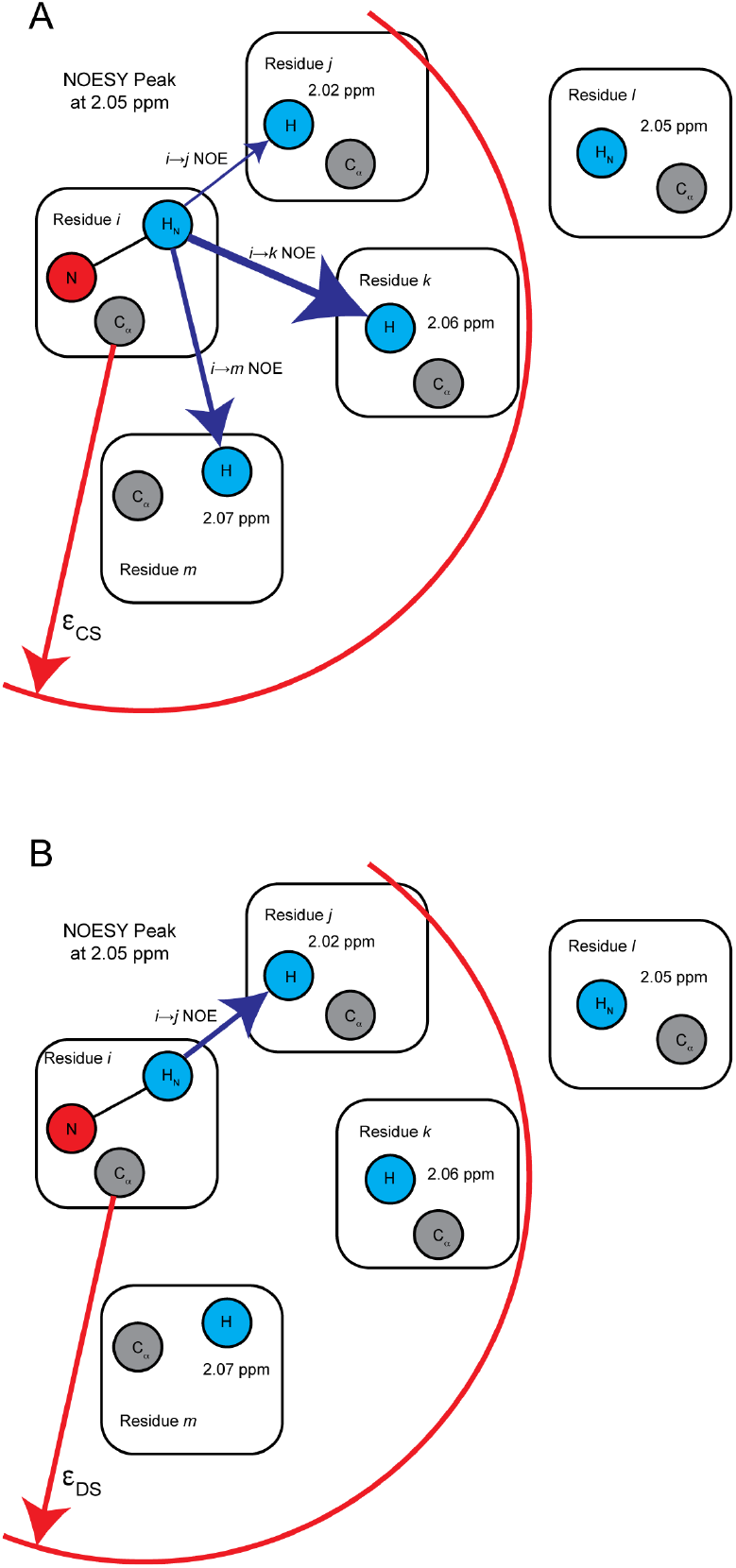
Calculation of Contact and Distance Scores. The HN of residue *i* is seen to have a NOESY cross peak at a shift of 2.05 ppm. All potential contacts with a residue separation of 4 or more are shown as residues *j,k,l*, and *m*. A. The CS is calculated by considering all residues whose C_*α*_ distance is less than the cutoff *ϵ*_*CS*_. In the final CS, the *ik* NOE will contribute the most as its chemical shift agreement is best, as indicated by the bolder blue arrow. B. In the DS, only the potential contact with the shortest HN to H distance is considered, indicated by the blue arrow between *i* and *j*. If the interresidue distance is greater than *ϵ*_*DS*_, it is counted towards the final DS.

The scores for individual interresidue contacts are summed and then normalized by the maximum number of contacts with a separation of at least five residues, given by (*n*_*resi*_ − 4)(*n*_*resi*_ − 5), where *n*_*resi*_ is the number of residues compared. CS has been designed to be high when the structure and data are good matches. In this case, most of the off-diagonal peaks in the NOESY spectrum due to long-range contacts are assigned to contacts seen in the proposed structure and contribute to CS. Higher confidence in the assignments and more observed NOESY crosspeaks lead to a higher value of this score. Therefore, a high CS should correspond to a good structure with a large number of reliably assigned NOESY crosspeaks.

#### 2.2.3 Distance Score

The calculation of DS is illustrated in Figure 3B and is detailed in Algorithm 3. DS is the fraction of NOESY crosspeaks that cannot be assigned by the proposed structure. To calculate DS, all potential contacts for a single peak are reduced to the single interresidue contact that has the smallest separation according to the proposed structure. If this separation is greater than the threshold *ϵ*_*D*_, the peak is counted in the DS score. This indicates that that peak cannot be reasonably explained by the proposed structure. As for CS, only long-range contacts are considered. The final DS is the ratio of crosspeaks that cannot be explained by the proposed structure to the total number of cross peaks. The closer DS is to 1, the worse the agreement between the NMR data and the proposed structure.

The threshold *ϵ*_*D*_ is selected separately from the contact distance threshold *ϵ*_*CS*_ used in CS. *ϵ*_*CS*_ is the maximum distance between the *α* carbons for two residues to be considered ‘in contact’ while *ϵ*_*D*_ is the maximum C*α* distance between two residues that can give rise to an observable NOESY peak. *ϵ*_*D*_ was determined via a grid search to optimize the correlation between the DS and TM-score, as detailed in the SI. It was found to be 10 Å.

Filtering the potential contacts to a single interresidue contact distinguishes DS from CS. CS considers all potential contacts weighed by their assignment certainty, while DS reduces each peak to a single potential contact that best represents the peak.Because of the r^*−*6^ dependence of the NOE intensity, the chosen potential contact used in DS is expected to make the primary contribution to the cross peak.

## 3 Results

### 3.1 Testing CS and DS on example structures: Aq1974 and pro-IL-18

To highlight the usefulness of CS and DS, we discuss their application to two example proteins, both of which are known to be difficult to model even with AlphaFold, but their structures have been determined by NMR spectroscopy. By comparing the values of CS and DS on proteins of known difficulty for AlphaFold prediction, we can begin to gain some insight into their value.

Aq1974 (PDB ID: 5SYQ) is a small protein of unknown function (puf) from *Aquifex aeolicus* with an unusual amino acid sequence that has an exceptionally high aromatic content, 14% of the folded region of the protein. Due to its unusual primary sequence, the CASP-10 competition found that the structure of Aq1974 is difficult to predict.[7] It is also challenging for AlphaFold2. AlphaFold2’s internal reliability testing algorithms(MSA coverage, alignment error, and LDDT) all warn that this prediction is probably not reliable. Another protein that has been found to be a difficult target for AlphaFold2 is Pro-interleukin-18 (pro-IL-18, PDB ID: 8urv), which is the precursor form of the cytokine IL-18. Kay and coworkers have shown that its 3D structure as determined by NMR disagrees significantly from that predicted by AlphaFold2[6]. We use these structures to test our new heuristics.

To examine the scaling of CS and DS with prediction accuracy, we chose one of the NMR models of both Aq1974 and pro-IL-18 as their reference model. TM-scores were then calculated with respect to these reference models. The average value of the TM-score in the NMR structural ensemble for Aq1974 is 0.860 ± 0.037 while that for pro-IL-18 is 0.905 ± 0.010. In the AlphaFold2 prediction of the structure of Aq1974, there are two classes in the top five models. The first class, colored orange in Figure 4, contains two members and gave an average TM-score of 0.587 ± 0.005. The remaining 3 predicted structures, colored red, gave an average TM-score of 0.327 ± 0.017. AlphaFold2 performed equally poorly for all its predictions of pro-IL-18, where the average TM-score with the reference structure of the five predicted structures is 0.510 ± 0.001.

**Fig. 4:**
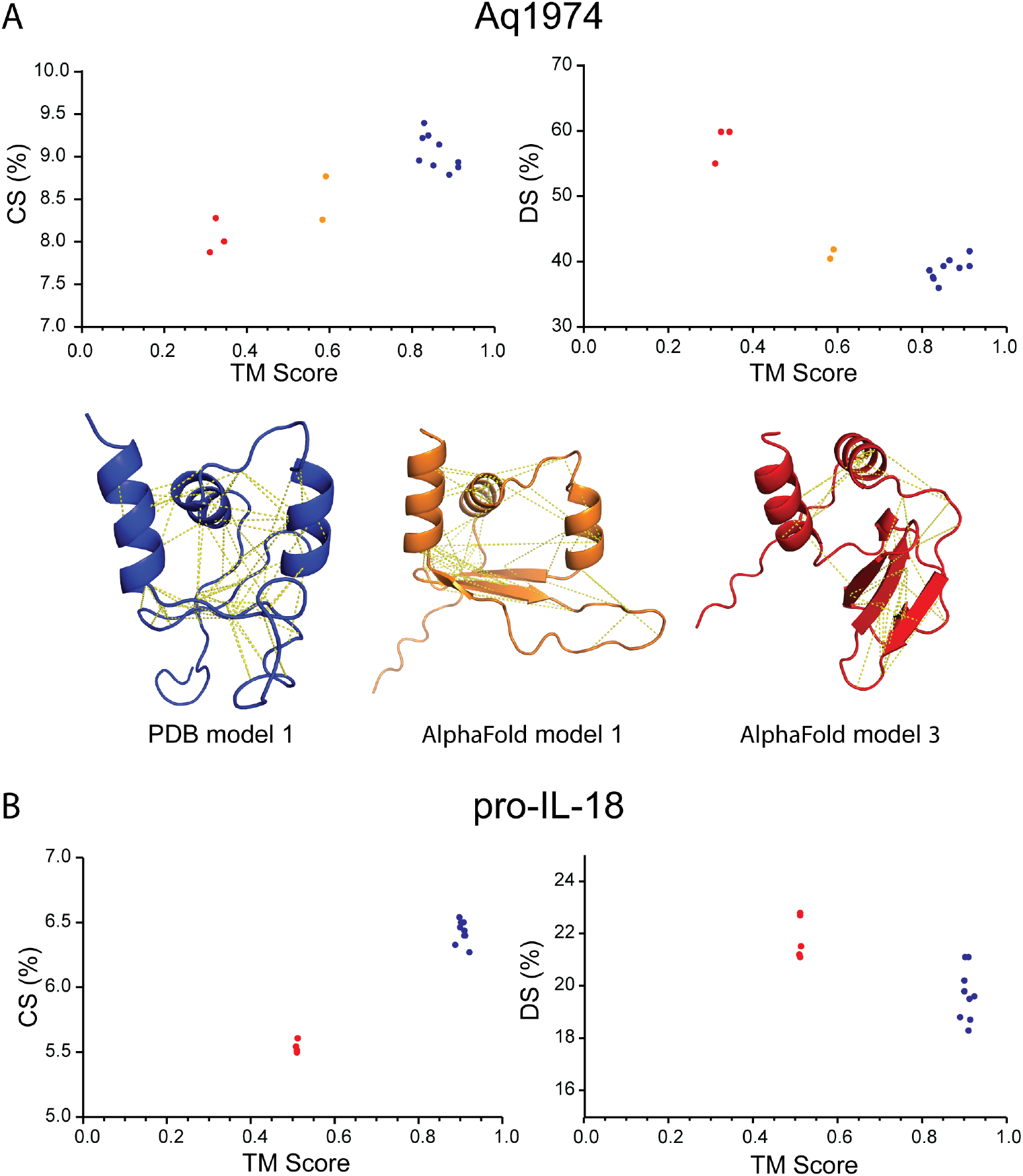
Contact Score (CS) and Distance Score (DS) as functions of TM-score for Aq1974 and pro-IL-18. A) CS and DS as a function of TM-score for Aq1974. Blue points are from the PDB structure where the first model structure was used a the reference model. Also included are the first model from the PDB structure (blue) and two AlphaFold models: orange was the best alpha fold model while while the red models is an example where the TM-score was about 0.35. The dashed yellow lines are the assigned long range interresidue connections seen in the NMR data and identified by comparing with the structure. Red and orange points correspond to red and orange models. B) CS and DS as a function of TM-score for pro-IL-18. AlphaFold2 only predicted one class of incorrect models labeled red.

Figure 4 shows example models of Aq1974: NMR determined structure colored blue, the best AlphaFold model colored orange, and an example of the second group of poor AlphaFold models colored red. The dashed yellow lines on the structural models indicate the identified long-range interresidue contacts from the comparison of the NMR data with the proposed structure. The number of identified contacts decreases as the quality of the model decreases. This correlates directly with CS and DS. For both Aq1974 and pro-IL-18 show monotonic relationships between CS, DS, and TM. CS increases with increasing values of TM while DS decreases in both of these structures. This indicates that these scores are useful in comparing structures to NMR data.

### 3.2 Testing Heuristics on the Compiled Dataset

Although examples like Aq1974 and pro-IL-18 demonstrate that CS and DS correlate with TM-score for these cases, we need to demonstrate that they are useful in a broader context. CS and DS were calculated over the entire compiled dataset of 3085 structures described in section 2.1. To increase the number of poorly predicted structures, the structure and NMR data of each member of the original dataset were associated with an AlphaFold prediction taken from another randomly chosen member of the dataset. These randomly chosen predictions are usually quite different from the experimentally determined structure, increasing the number of poorly predicted structures. We call the dataset that includes these ‘mismatched’ structures the ‘augmented dataset’ and it contains 6170 entries. It provides a useful way of examining the behavior of CS and DS on a wider variety of predicted structures than AlphaFold could typically provide.

Figure 5A shows CS and DS as a function of TM-score on both the original and mismatched datasets. The TM-score was calculated between the AlphaFold structure and the first model of the NMR ensemble. Each point in the scatter plots is colored by whether the sample comes from the mismatched (red) or the original (blue) data. The augmented dataset is the union of these two sets. This figure shows that in the original dataset contains more ‘good’ structures, while the mismatched experiment introduces more ‘poor’ structures, as indicated by the shift of the distribution between the blue and red points. This is expected from the success of AlphaFold; however, there are still a significant number of structures in the original dataset (blue) whose TM-scores are less than 0.5, indicating problematic predictions. While the observed correlation between CS and DS with the TM-score is noisy in Figure 5, it is clearly present. We further tested CS and DS on the 117 members of our data set that contained NOESY peak lists, Figure 5B. The correlations seen in the augmented dataset are clearly visible in data with experimental NOESY peaks lists.

**Fig. 5:**
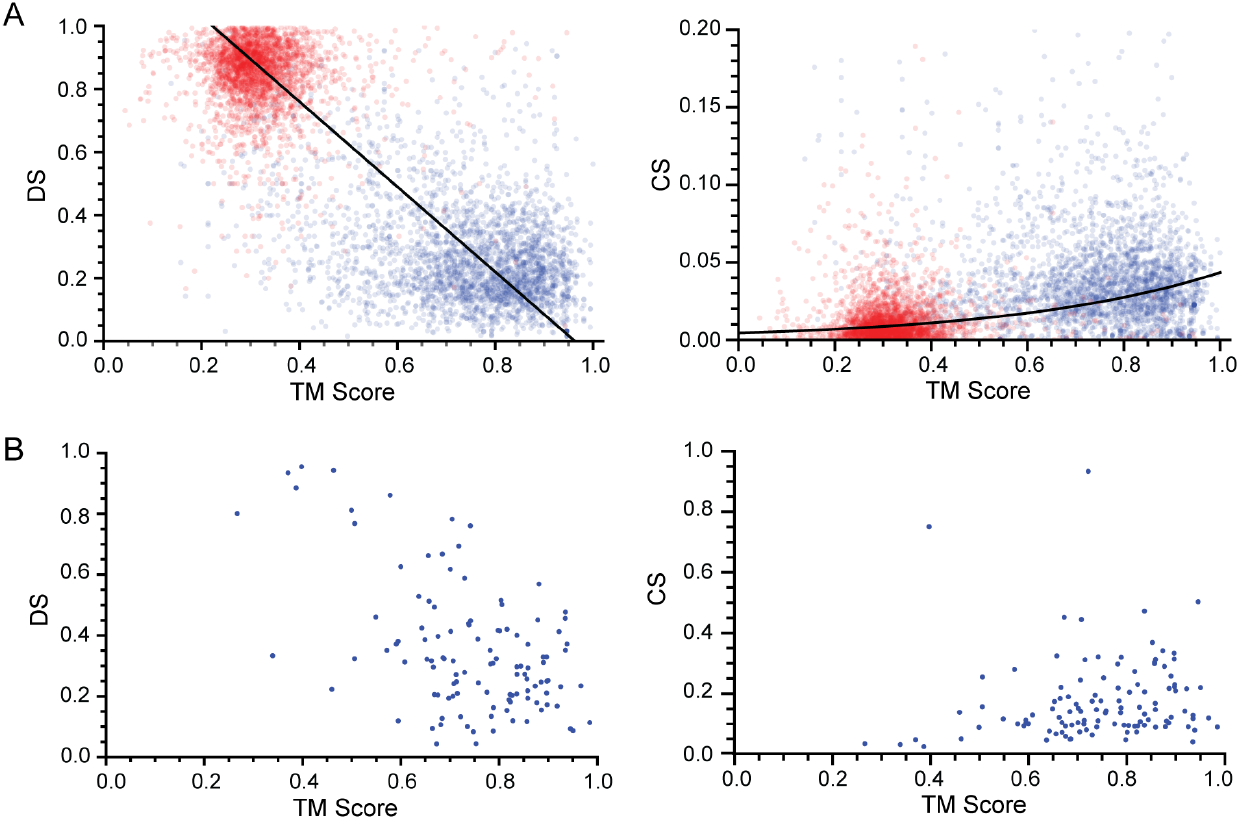
CS and DS as a function of the TM-score. A) Blue data are from the original dataset, while red are from intentionally mismatched data. The number of structures in a region is indicated by the shade of red and blue: the darker the color, the more data represented. While there is considerable scatter in the data, it is clear that there is a positive correlation between CS and TM-score and a negative correlation between DS and TM-score over the entire dataset, indicated by the black curves. B) Correlaton of CS and DS with TM-score over the 117 members of our data set that contained NOESY peaklists.

To quantify the correlations between these heuristics and our datasets, Spearman’s rank correlation coefficients were calculated between all input parameters and the TM-score, Figure 6. Figure 6A shows the correlation coeffients for the original and augmented datasets, upper and lower triangles, respectively, while Figure 6B shows that for the dataset with experimental NOESY peaks lists. Spearman’s rank correlation coefficients are a nonparametric method of determining whether two parameters are monotonically dependent upon each other. The more typically used Pearson’s correlation coefficients assume that the parameters are linearly related and the data are Gaussian distributed, both of which are not necessarily true for our dataset. These assumptions are removed when using Spearman’s rank correlations.[29] These correlation data show the power of using the augmented dataset. For example, CS shows a small but statistically insignificant correlation with TM-score in the original dataset; however, upon adding mismatched data, this correlation becomes positive and highly statistically significant, *p <<* 0.0001. The addition of mismatched structures increases our confidence in correlations between our new heuristics and the TM-score by increasing the number of poor structures in the dataset. Figure 6B shows the correlation coefficients over the dataset with experimental NOESY peak lists. In the case with experimental data, PAE, CS, and DS are all significantly correlated, *p <* 0.01, with the TM-score as also seen with the augmented data in Figure 6A. This confirms the results seen in the augmented dataset. We conclude that these heurisics measure important characteristics of the agreement between predicted structure and “true” structure and allows us to use them as basis for characterizing predicted structures.

**Fig. 6:**
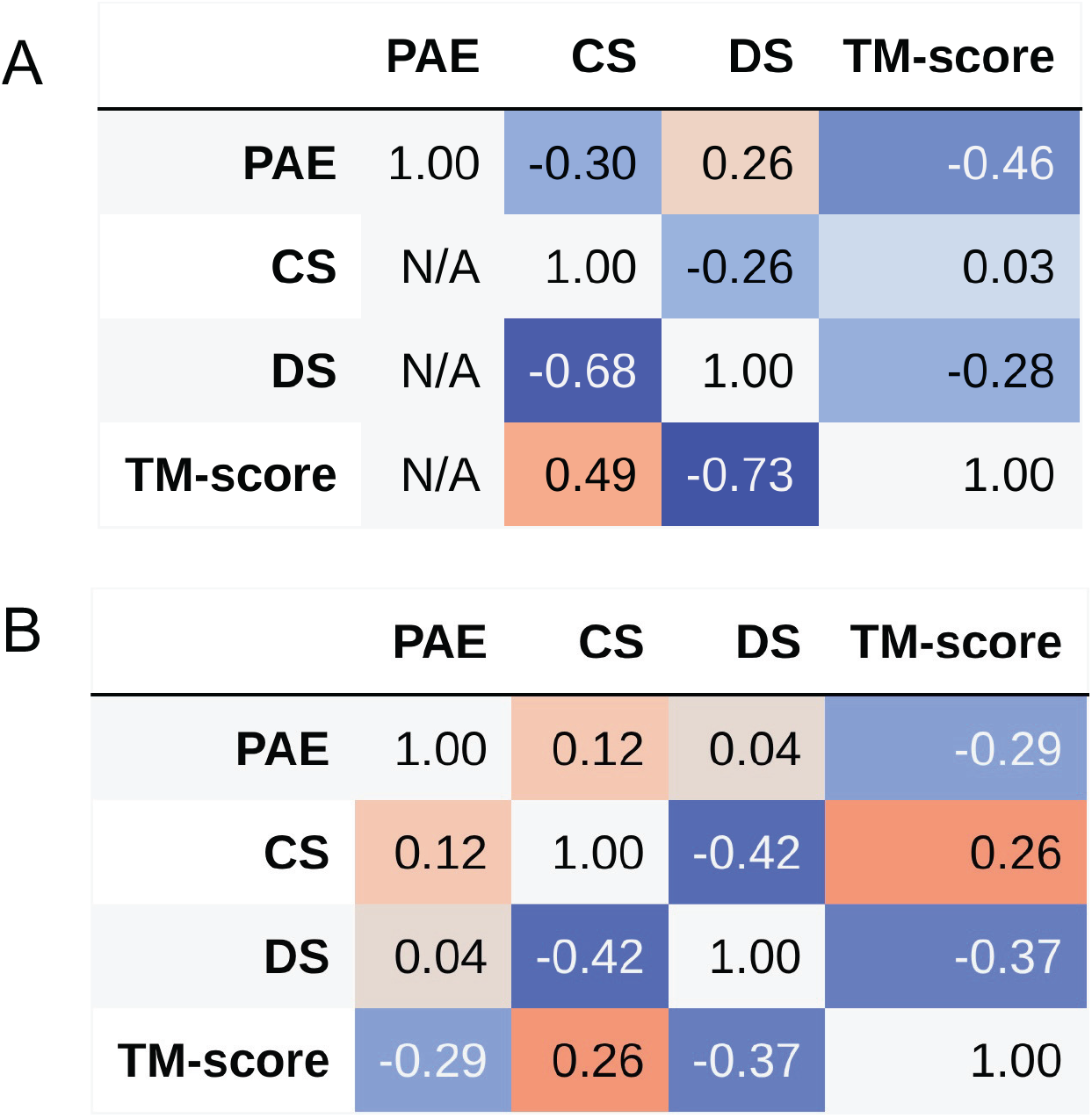
Spearman’s rank correlations between heuristics and TM-score. Large positive values are in red and large negative values are in blue. A) Correlations presented in the upper triangle are from the original dataset, while the lower triangle comes from the dataset augmented with mismatched experiments. The final columns and the final row give the correlations with the TM-score from the regular experiments and the mismatched experiments, the ground truth measure of quality in each experiment. The predicted aligned error (PAE) and distance heuristic have the highest correlations with the ground truth in the regular experiment, while both of our heuristics have a strong correlation in the mismatched experiment. B) Correlations in the dataset with experimental NOESY peaks lists.

### 3.3 Structural Prediction Assessment by NMR (SPANR)

We developed a small machine learning method, specifically a support vector machine (SVM), to predict the quality of the AlphaFold structure based on our heuristics. This is shown in the left two columns of Figure 1. An SVM is a simple machine learning algorithm that, in this case, we have trained to place AlphaFold predicted structures into one of two categories: consistent or inconsistent with the NMR data. We call our SVM Structural Prediction Assessment by NMR (SPANR).

#### 3.3.1 Training

Developing multiple SVMs allows us to determine our heuristics’ ability to experimentally test the consistency of AlphaFold2 predicted structures with experimentally determined data. Predicted structures were classified as “consistent” if the TM-score between predicted and experimentally determined structures was greater than 0.5. Multiple metrics were used as inputs to the SVMs which were split into groupings of either PAE or CS-DS. We trained three SVMs using one or both of the groupings on the original dataset, see Table 1, allowing us to determine the effect of each combination on the accuracy of the SVM to categorize the predicted structure. To perform the training, we split each dataset into training and test sets with an 80-20 split, then used scikit-learn to fit the SVM on the training data [30]. We also trained an SVM on the augmented dataset using only the CS and DS as inputs.

**Table 1:**
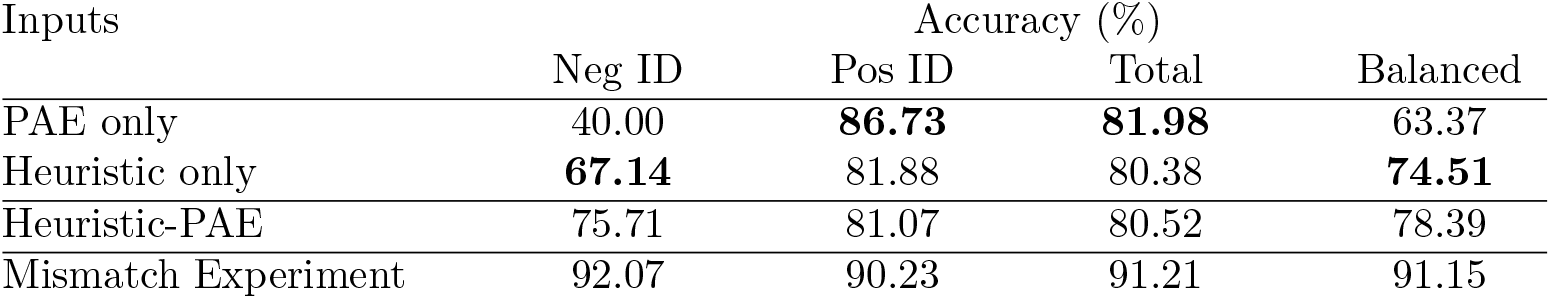
Comparison of accuracy of SVMs with different inputs on the binary classification task of identifying accurate (TM-score *>* 0.5) structures. Negative and Positive ID rates refer to the percent of negative or positive structures correctly identified as such by the SVM. The SVMs were all trained with balanced class weights, therefore we include the balanced accuracy to demonstrate how well each model has achieved the goal of equally identifying positive and negative samples. Models above the line use only one class of inputs, while those below are combinations of inputs. Finally, the results from the mismatch experiment are included, in which the SVM was given only the heuristics as inputs.

#### 3.3.2 Accuracy

Each of the three SVMs were trained on the original dataset and scored on the test set. Due to presence of many more well-predicted structures in the original dataset, we consider the balanced accuracy to be the best metric for comparing SVMs. We are particularly interested in each classifier’s ability to identify models inconsistent with the NMR data, because false negatives are more tolerable in most structure determination cases; *i.e*., verifying a well-predicted structure is preferable to treating a poorly predicted structure as correct. The results are collected in Table 1. Of the two single group inputs, CS-DS perform the best at identifying inconsistent structures. PAE performs best at identifying consistent structures, succeeding 87% on the test dataset, but only succeeding 40% in predicting inconsistent structures. This suggests that the predicted aligned error overestimates AlphaFold’s accuracy. CS and DS are capable of correcting these inaccuracies. The rate of identifying consistent structures dropped to 82% using just the DS and CS scores, but the rate of identifying poor structures increased to 67%. Using all inputs together gives the best results, resulting in a balanced accuracy of 78%, with consistent structures identified at a rate of 81% and inconsistent structures at a rate of 75% in the test dataset.

Sci-kit learn has the option to predict the confidence of each class categorized by the SVM, rather than predicting the single most likely class. To calculate the confidence of the classes, a slightly different model is used than in the baseline SVM. Figure 7 shows the TM-score for every member of the test dataset as a function of the classification confidence. Figure 7A uses the CS-DS-PAE SVM over test data from the original dataset while Figure 7B shows the results from the CS-DS SVM on the test data from the augmented dataset. The augmented data cannot use PAE as input due to the inclusion of mismatched data. The red dotted lines indicate predictions above 60% confidence in either class, consistent or inconsistent; greater than the red line at 0.6 indicates greater than 60% confidence that the predicted structure is consistent with the NMR data and less than that at 0.4 means greater than 60% confidence that the structure is inconsistent. The SVMs show very high accuracy in these regions with improvements to above 90% for inconsistent results using the augmented data. In fact, when the model is over 80% confident in a positive sample, it is 94% accurate. If the SVM is 40% to 60% confident, the model is only 50%-60% accurate. Figure 7C tests the CS-DS-PAE SVM over the dataset with experimental NOESY peak lists. This shows that with experimental NOESY data, this SVM has an accuracy of over 95% when the confidence is above 60% and is about 80% accurate in the confidence range 40% to 60%. Unfortunately, due to the small size of the dataset with experimental NOESY spectra, no examples were found in the confidence range between 0% and 40%.

**Fig. 7:**
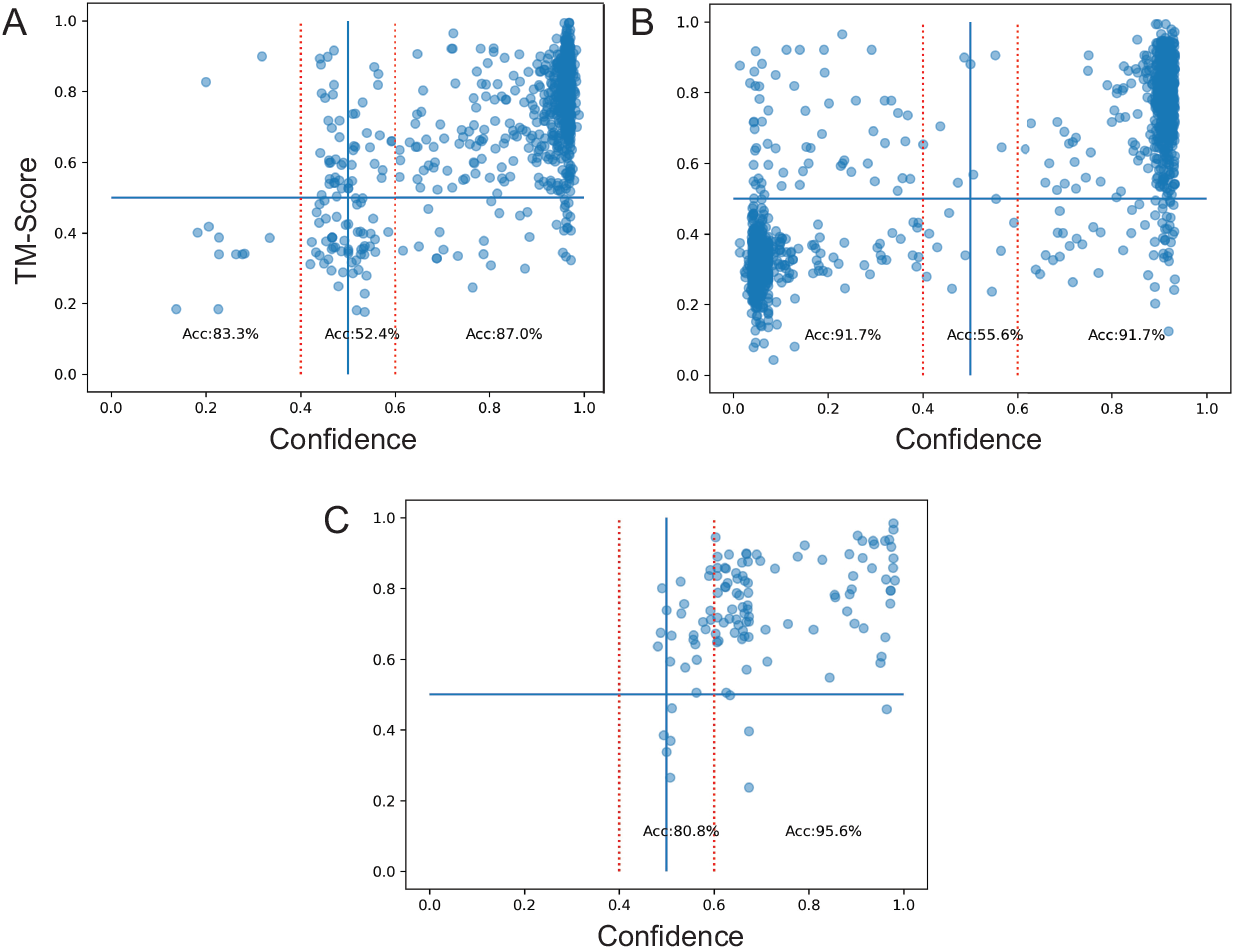
TM-score as a function of the confidence the data are consistent this the predicted model from our SVMs. Blue dots are TM-Score/confidence pair for each structural prediction over the test data set. The horizontal blue line indicates a TM-score of 0.5 which is the value we chose for a good prediction of the true structure. The vertical red lines are at 0.4 and 0.6 confidence levels. A) SVM trained and calculated on the original dataset. B) Trained and calculated on the augmented dataset. C) SVM tested on the experimental dataset.

To test our SVMs ability to detect inconsistent structures, the CS-DS SVM trained on the original dataset was applied to a test set taken from just the mismatched data and found to have a total accuracy of 95%. A new CS-DS SVM trained on the augmented dataset using the same train-test split then analyzed the same test set from the mismatched data and was found to have an accuracy of 94%. Thus, training on the augmented dataset did not improve the SVMs ability to identify inconsistent models over the original SVM using the dataset without the mismatched data. We conclude that while mismatched samples are important to verify correct performance of the CS and DS on very poor samples, they do not help the SVM learn to separate consistent from inconsistent structures more than the original dataset.

### 3.4 Applications of SPANR

To demonstrate the power of SPANR, predicted structures and NMR data of two known structures, prot-IL-18 and Aq1974 which are not part of our training or test sets, were analyzed by SPANR. Additionally, we examined a designed protein, LoTOP, whose structure has not been previously determined with SPANR. These examples demonstrate the power of our SVM.

#### 3.4.1 pro-IL-18 and Aq1974

SPANR analyzed the consistency of the predicted structures with NMR data of the proteins pro-IL-18 and Aq1974 discussed above. Using the best AlphaFold2 model for pro-IL-18 and entering its NOESY peak data, chemical shift assignments, and AlphaFold’s PAE file, SPANR finds that this structure is consistent with the NMR data with a 93% certainty. This means that with 93% certainty the TM-score between this structure and the “true” structure is greater than 0.5. The TM-score between this predicted structure and model 1 of the NMR determined structure was found to be 0.51.

In the case of Aq1974, we can examine predicted structures from both classes shown in Figure 4. We find that SPANR identifies AlphaFold model 1 as consistent with a confidence of 88% and that of model 3 as inconsistent with a confidence of 69%. The TM-scores of the first model of the NMR determined ensemble to the first and third predicted structures were 0.58 and 0.31, respectively.

#### 3.4.2 LoTOP

LoTOP is a fast folding variant of Top7 that was designed by permuting the secondary structure elements of Top7. Circular permutation is a typical method for creating a new version of an existing protein. This method alters the protein’s topology by linking the N and C termini and nicking one of the existing loops; unfortunately, the N and C termini are too far apart in Top7 to apply this method. Hence, two new archetypical beta-turn sequences[31] were introduced into Top7 to rewire its secondary structures into a new topology.[32] Instead of Top7’s (*β*)_2_-H-*β*-H-(*β*)_2_ order, the permutant “LoTOP” has a H-(*β*)_5_-H order, a simpler topology with a significantly lower relative contact order (RCO) (SI Figure 4). Although LoTOP is not strictly a new fold (e.g., bacterial, yeast and human Frataxins have the same gross fold but are H-(*β*)_7_-H), LoTOP can be considered a new superfamily. Our first design attempt was successful. LoTOP retains Top7’s high stability (12 kcal mol^*−*1^, SI Figure 5).

^1^H^15^N HSQC spectra of LoTOP gave a high-quality spectrum with most HN peaks observable (SI Figure 6), indicating a stable folded structure. The spectrum was assigned by routine methods, resulting in 97% of the backbone shifts and 80% of all ^1^Hs assigned. The product map between the potential contact map of possible assignments of the NOESY peaks and the AlphaFold predicted contact map for the structure was calculated. Figure 8A shows this product map for the highest confidence AlphaFold2 structure, where the color scale represents the score for a long-range inter-residue contact as defined in section 2.2.2. CS is the sum of the off-diagonal elements of this matrix, shown as the yellow region of the map, normalized to the area of this region. The average CS over the 5 best AlphaFold structures is 0.117±0.001 while the average DS is 0.316±0.002. Comparing these values with the distribution of the scores over the augmented dataset, Figure 4, the TM-scores to the “true” structure is expected to be greater than 0.8, indicating that the AlphaFold structures are a good representation of the “true” structure. These data were input into our SVM that uses CS, DS and PAE and trained on the original dataset, where it finds that the NMR data are consistent with the proposed structure with a confidence of 97%.

**Fig. 8:**
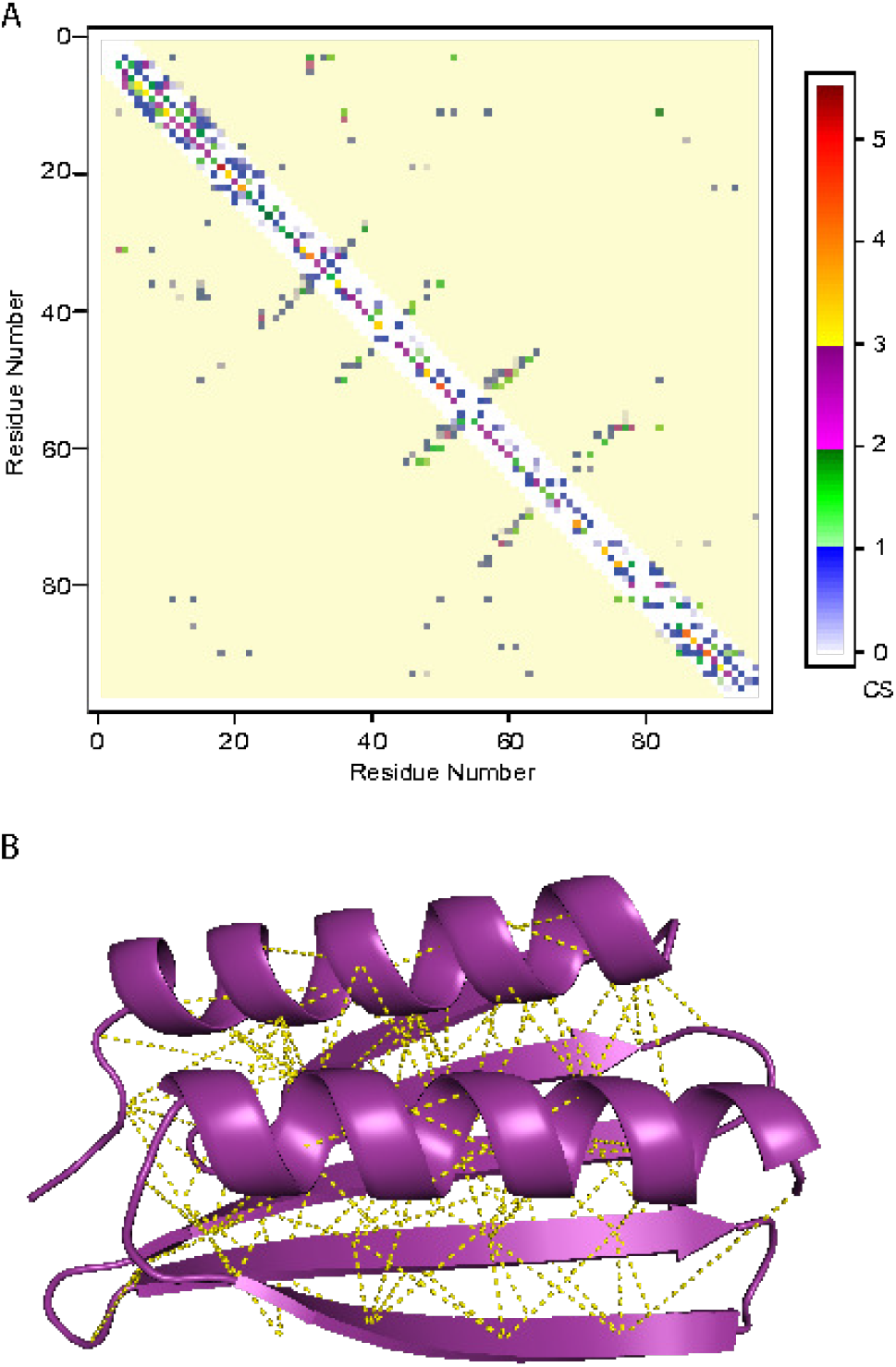
Agreement between NMR data and AlphaFold2 predicted structure of LoTOP. A.Product of the potential contact map and the AlphaFold2 predicted contact map. Sum of the long range off-diagonal elements of this map (yellow region) gives the CS. B.The AlphaFold2 predicted structure with the NMR determined long range contacts.

## 4 Discussion

These results demonstrate the power of simple NMR experiments in testing AI-derived structural models. By measuring a limited set of NOESY constraints, just from the ^15^N-edited NOESY experiment, SPANR can distinguish structural models that are consistent or inconsistent with the data. To train SPANR, we developed CS and DS to quantify the agreement between the NMR data and the predicted structure.

CS and DS are unique in that they quantify the agreement between structure and NMR data by analyzing interresidue contacts rather than specific H-H contacts. This abstraction allows one to forgo detailed assignment of ^1^H resonances to atom positions, but rather only identify from which residues the ^1^H peaks arise. Despite this course-graining of of NOESY interactions, both CS and DS are highly correlated to the TM-score between the NMR-determined and the predicted structure. Examining CS in more detail, we did find that its value is dependent upon the chemical shift uncertainty,Δ_*shift*_ used in its calculation, making it a reliable measure of agreement between NMR data and structure for a single sequence, but noisy over a large dataset where different chemical shift uncertainties are expected. We have shown that our SVM has higher accuracy when trained with CS, demonstrating its usefulness. DS does not suffer as much from this dependence on chemical shift uncertainty. Of particular interest, CS and DS do not require full assignment of all H peaks to individual atoms but only to residues because these heuristics only look at interresidue contacts. This reduces the work required to use CS and DS. These quick and simple methods validate AlphaFold structures, and will serve as the basis for more complex validation and refinement methods.

CS and DS are valuable as inputs into SVMs to determine the consistency of NMR data with proposed structures. SPANR has been shown to be useful in performing this categorization. When SPANR determines that the certainty of these categories is above 70-80%, the categorization accuracy is well above 90%. This allows us to use NMR data to decide if further refinement of the proposed structure is necessary.

The results in this paper rely on the large dataset of NMR determined and AlphaFold2 predicted structures, and the corresponding NMR data that we compiled. This dataset will be key in the development of novel tools to accelerate protein structure determination through the use of hybrid AI-experiment methods; however, the dataset needs improvement, which will occur by increasing both its size and the quality of its structural constraints. One improvement we made to this dataset was to include intentionally mismatched data. This significantly increased the number of poorly predicted structures in the augmented dataset produced. One major issue with the compiled dataset is its lack of NOESY constraints for most of its members. Due to this limitation, we had to calculate the expected NOESY peaks from the true structure and the assigned chemical shifts. An ideal dataset would include a comprehensive set of experimentally determined NMR structural constraints which could be used to train hybrid AI-experimental models. The main bottleneck to this is finding reported experimental NMR results, for which the BMRB is the largest source, but not necessarily the only. Besides finding other sources of data, adding new experimental results to the dataset as they are obtained will be important. The BioNMR community is developing tools that will help homogenize and collect newly captured experimental data [33] which will be crucial in this.

Ultimately, we hope that the recent advances in AI and machine-learning will increase our ability to solve structures by NMR quickly and accurately. The future of structural NMR will be refining predicted structures based upon NMR constraints. Klukowski et al. have shown that using AI derived structures can increase the efficiency of assigning protein NMR spectra. Ultimately, the goal is to glean biological information in the most efficient way possible, by using hybrid methods that combine AI with experiment.

## 5 Methods

### 5.1 LoTOP Expression and Purification

A single colony of E. coli BL21(DE3) C+ that carries the recombinant plasmid vector pET-21-8xHis-DesG-LoTOP was inoculated in 5 mL LB broth containing 1:1000 (Ampicillin 100µg/mL) overnight at 37°C on shaker at 225 rpm. This culture was added into 1L of M9 minimal medium containing 100ug/mL Ampicillin and supplemented with ^15^NH_4_Cl (1g/ L) and incubated at 37°C until an OD600 of 0.6-0.8 was reached. The temperature was then lowered to 20°C, and the flask was cooled for 30 minutes. Following this, protein induction was carried out with 1mM IPTG, and the culture was incubated overnight at 225 rpm. The next morning, the induced expression culture was harvested by centrifugation at 5500rpm for 15 minutes at 4°C and the cell pellet frozen at -80°C.

The frozen cell pellet was resuspended in 50mM Tris-HCl pH 7.5, 150mM NaCl, containing a cocktail of protease inhibitors (Abcam ab27005). The cells were then lysed by sonication (Branson Sonifier 450) on ice. The cell debris was removed by centrifugation at 12,500 rpm, and 4°C for 30 minutes. The protein was then purified using metal affinity chromatography. The binding and washing steps were carried out in 50mM Tris-HCl pH 7.5, 150mM NaCl, and 20mM imidazole, and the protein was eluted with 250mM imidazole. The His-tag was removed by incubating overnight at 4°C with 6xHis-TEV (Tobacco Etch Virus) protease. Metal affinity chromatography was then used to remove the 8xHis-DesG tag and 6xHis-TEV protease from the solution. The protein was buffer exchange for long-term storage using a Superdex 200 column into 10mM Phosphate buffer at pH 7.0. Finally, for NMR studies, the buffer was exchanged, when needed, into different buffers depending on the pH using a G-25 Sephadex spin column.

### 5.2 NMR Spectroscopy

NMR spectra were acquired on Bruker AVANCE IIIHD 600 MHz NMR spectrometer equipped with a room temperature TXI probe. Spectra were acquired at a sample temperature of 37^*°*^ C in 20 mM MES and 50 mM NaCl buffer at a pH 6.5. Assignments of U-[^13^C, ^15^N]-LoTOP were obtained by analysis of standard triple resonance NMR spectra. This resulted in the assignment of 96.8% of the backbone atoms and 79.6% of all ^1^H atoms.

## Acknowledgements

JW was supported with funding by the University of Chicago Data Science Institute (DSI). IAG and the development of LoTOP were supported by NIH grant GM055694. The authors thank James R. Hinshaw, Tiffany M. King, and Tobin R. Sosnick for designing and characterizing LoTOP and Michael Baxa and Heewhan Shin for valuable conversations. All NMR data were acquired in the Biomolecular NMR Core at the University of Chicago.

## Data Availability

Code and data references will be made available at the time of publication via GitHub. Structures and data downloaded from the PDB (https://www.wwpdb.org), BMRB (https://bmrb.io/), and AlphaFold DB (https://alphafold.ebi.ac.uk).

## A Algorithms

### Algorithm 1

NOESY Simulation

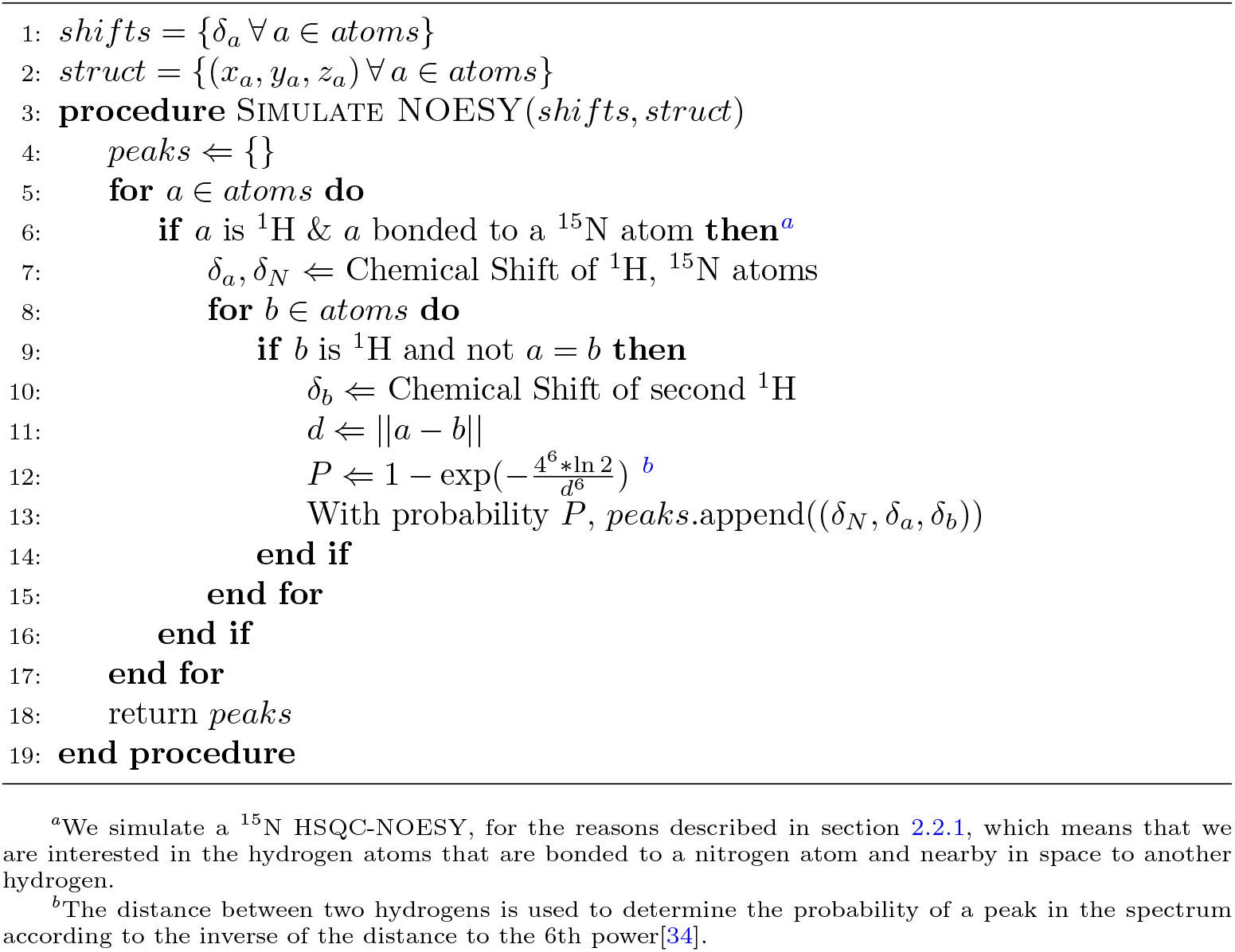

### Algorithm 2

Contact Score

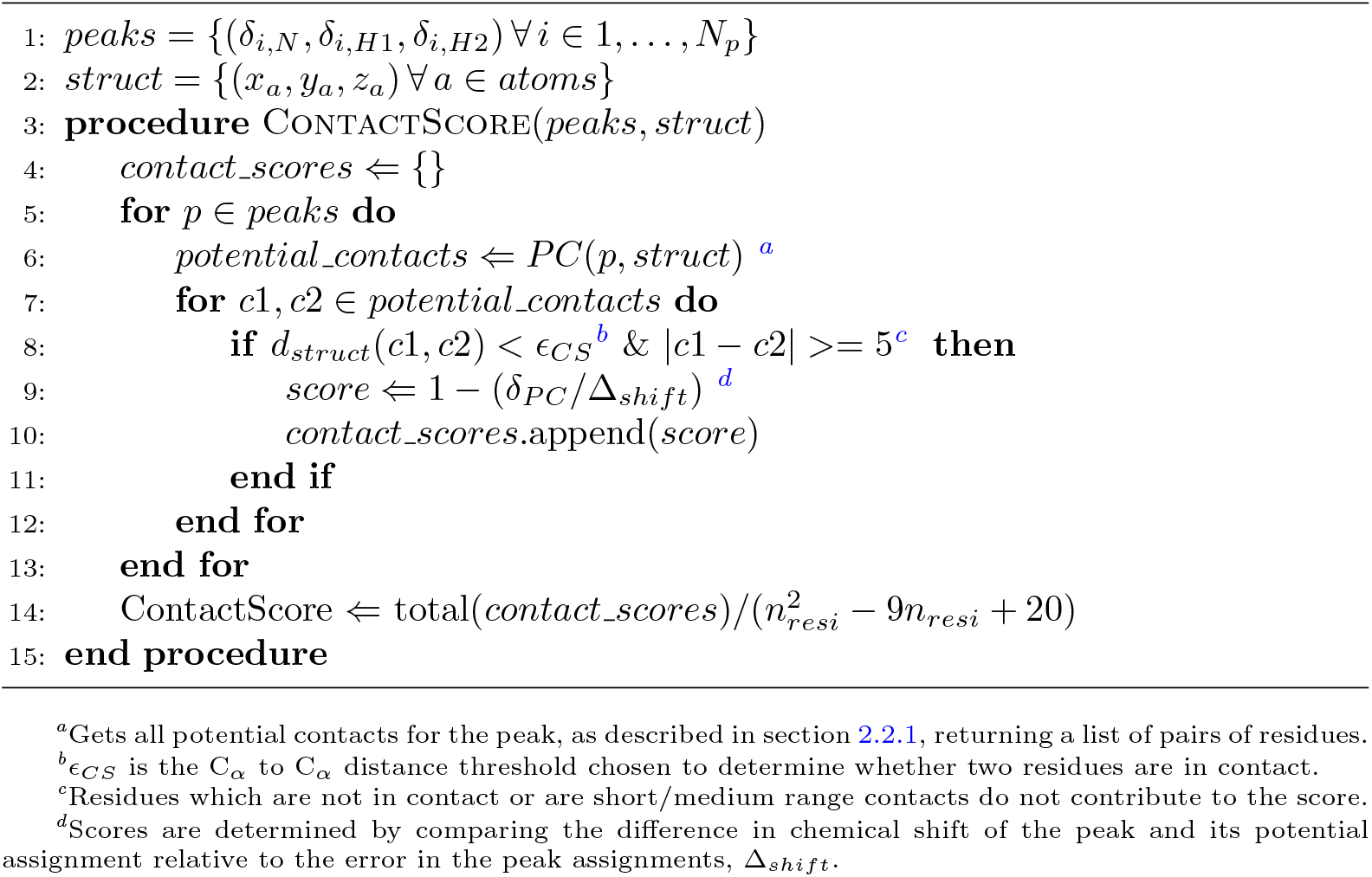

### Algorithm 3

Distance Score

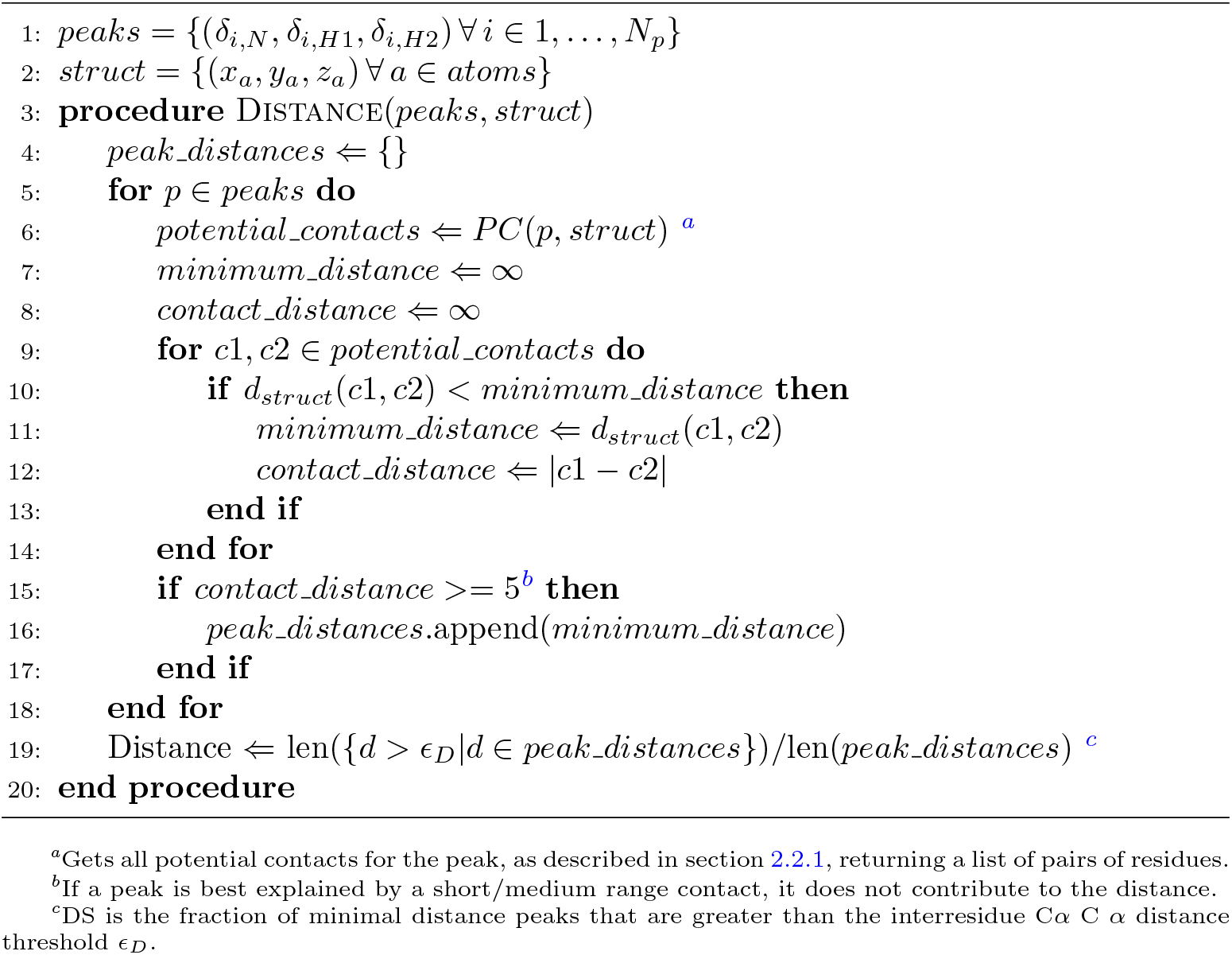

## References

[1] Anfinsen, C.B.: Principles that govern the folding of protein chains. Science 181(4096), 223–230 (1973) 10.1126/science.181.4096.223 https://www.science.org/doi/pdf/10.1126/science.181.4096.223

[2] Dill, K.A., Ozkan, S.B., Shell, M.S., Weikl, T.R.: The protein folding problem. Annual Review of Biophysics 37([Volume 37, 2008, Volume 37,]), 289–316 (2008) 10.1146/annurev.biophys.37.092707.153558

[3] Jumper, J., Evans, R., Pritzel, A., Green, T., Figurnov, M., Ronneberger, O., Tunyasuvunakool, K., Bates, R., Žídek, A., Potapenko, A., Bridgland, A., Meyer, C., Kohl, S., Ballard, A., Cowie, A., Romera-Paredes, B., Nikolov, S., Jain, R., Adler, J., Hassabis, D.: Highly accurate protein structure prediction with alphafold. Nature 596, 1–11 (2021) 10.1038/s41586-021-03819-2

[4] Kryshtafovych, A., Schwede, T., Topf, M., Fidelis, K., Moult, J.: Critical assessment of methods of protein structure prediction (casp)—round xiv. Proteins: Structure, Function, and Bioinformatics 89(12), 1607–1617 (2021) 10.1002/prot.26237 https://onlinelibrary.wiley.com/doi/pdf/10.1002/prot.26237

[5] Buel, G., Walters, K.: Can alphafold2 predict the impact of missense mutations on structure? Nature Structural & Molecular Biology 29, 1–2 (2022) 10.1038/s41594-021-00714-2

[6] Bonin, J.P., Aramini, J.M., Dong, Y., Wu, H., Kay, L.E.: Alphafold2 as a replacement for solution nmr structure determination of small proteins: Not so fast! Journal of Magnetic Resonance 364, 107725 (2024)

[7] Sachleben, J.R., Adhikari, A.N., Gawlak, G., Hoey, R.J., Liu, G., Joachimiak, A., Montelione, G.T., Sosnick, T., Koide, S.: Aromatic claw: A new fold with high aromatic content that evades structural prediction. Protien Science 26(2), 208–217 (2017)

[8] Tuinstra, R.L., Peterson, F.C., Kutlesa, S., Elgin, E.S., Kron, M.A.,, Volkman, B.F.: Interconversion between two unrelated protein folds in the lymphotactin native state. Proceedings of the National Academy of Sciences 105(13), 5057–5062 (2008)

[9] Ma, P., Li, D.-W., Brüschweiler, R.: Predicting protein flexibility with alphafold. Proteins: Structure, Function, and Bioinformatics 91(6), 847–855 (2023) 10.1002/prot.26471 https://onlinelibrary.wiley.com/doi/pdf/10.1002/prot.26471

[10] Bryant, P., Pozzati, G., Zhu, W., Shenoy, A., Kundrotas, P., Elofsson, A.: Predicting the structure of large protein complexes using alphafold and monte carlo tree search. Nature communications 13(1), 6028 (2022)

[11] Evans, R., O’Neill, M., Pritzel, A., Antropova, N., Senior, A., Green, T., Žídek, A., Bates, R., Blackwell, S., Yim, J., Ronneberger, O., Bodenstein, S., Zielinski, M., Bridgland, A., Potapenko, A., Cowie, A., Tunyasuvunakool, K., Jain, R., Clancy, E., Kohli, P., Jumper, J., Hassabis, D.: Protein complex prediction with alphafold-multimer. bioRxiv (2022) 10.1101/2021.10.04.463034 https://www.biorxiv.org/content/early/2022/03/10/2021.10.04.463034.full.pdf

[12] Abramson, J., Adler, J., Dunger, J., Evans, R., Green, T., Pritzel, A., Ronneberger, O., Willmore, L., Ballard, A.J., Bambrick, J., et al.: Accurate structure prediction of biomolecular interactions with alphafold 3. Nature, 1–3 (2024)

[13] Baek, M., DiMaio, F., Anishchenko, I., Dauparas, J., Ovchinnikov, S., Lee, G.R., Wang, J., Cong, Q., Kinch, L.N., Schaeffer, R.D., et al.: Accurate prediction of protein structures and interactions using a three-track neural network. Science 373(6557), 871–876 (2021)

[14] Tejero, R., Huang, Y.J., Ramelot, T.A., Montelione, G.T.: Alphafold models of small proteins rival the accuracy of solution nmr structures. Frontiers in Molecular Biosciences 9 (2022) 10.3389/fmolb.2022.877000

[15] Li, E.H., Spaman, L.E., Tejero, R., Huang, Y.J., Ramelot, T.A., Fraga, J.H. Keith J. and Prestegard Kennedy, M.A., Montelione, G.T.: Blind assessment of monomeric alphafold2 protein structure models with experimental nmr data. Journal of Magnetic Resonance 352 (2023) 10.1016/j.jmr.2023.107481

[16] Klukowski, P., Riek, R., Guntert, P.: Time-optimized protein nmr assignment with an integrative deep learning approach using alphafold and chemical shift prediction. Science Advances 9(47) (2023) 10.1126/sciadv.adi9323

[17] Tejero, R., Snyder, D., Mao, B., Aramini, J., Montelione, G.: Pdbstat: A universal restraint converter and restraint analysis software package for protein nmr. Journal of biomolecular NMR 56 (2013) 10.1007/s10858-013-9753-7

[18] Huang, Y.J., Powers, R., Montelione, G.T.: Protein nmr recall, precision, and f-measure scores (rpf scores): Structure quality assessment measures based on information retrieval statistics. Journal of the American Chemical Society 127(6), 1665–1674 (2005) 10.1021/ja047109h https://doi.org/10.1021/ja047109h. PMID: 15701001

[19] Fowler, N.J., Sljoka, A., Williamson, M.P.: A method for validating the accuracy of nmr protein structures. Nature communications 11(1), 6321 (2020)

[20] Lovell, S.C., Davis, I.W., Arendall III, W.B., Bakker, P.I.W., Word, J.M., Prisant, M.G., Richardson, J.S., Richardson, D.C.: Structure validation by cα geometry: ψ,ϕ and cβ deviation. Proteins: Structure, Function, and Bioinformatics 50(3), 437–450 (2003) 10.1002/prot.10286 https://onlinelibrary.wiley.com/doi/pdf/10.1002/prot.10286

[21] Hoch, J.C., Baskaran, K., Burr, H., Chin, J., Eghbalnia, H.R., Fujiwara, T., Gryk, M.R., Iwata, T., Kojima, C., Kurisu, G., Maziuk, D., Miyanoiri, Y., Wedell, J.R., Wilburn, C., Yao, H., Yokochi, M.: Biological Magnetic Resonance Data Bank. Nucleic Acids Research 51(D1), 368–376 (2022) 10.1093/nar/gkac1050

[22] Hall, S.R., Cook, A.P.F.: Star dictionary definition language: Initial specification. Journal of Chemical Information and Computer Sciences 35(5), 819–825 (1995) 10.1021/ci00027a005 https://doi.org/10.1021/ci00027a005

[23] Spadaccini, N., Hall, S.R.: Extensions to the star file syntax. Journal of Chemical Information and Modeling 52(8), 1901–1906 (2012) 10.1021/ci300074v https://doi.org/10.1021/ci300074v. PMID: 22725659

[24] wwPDB consortium: Protein Data Bank: the single global archive for 3D macro-molecular structure data. Nucleic Acids Research 47(D1), 520–528 (2018) 10.1093/nar/gky949

[25] Landrum, G.: RDKit: Open-source Cheminformatics. (2006). http://www.rdkit.org

[26] Ernst, R.R., Bodenhausen, G., Wokaun, A.: Principles of Nuclear Magnetic Resonance in One and Two Dimensions. Oxford Scientific Publications, ??? (1990)

[27] Varadi, M., Anyango, S., Deshpande, M., Nair, S., Natassia, C., Yordanova, G., Yuan, D., Stroe, O., Wood, G., Laydon, A., Žídek, A., Green, T., Tunyasuvunakool, K., Petersen, S., Jumper, J., Clancy, E., Green, R., Vora, A., Lutfi, M., Figurnov, M., Cowie, A., Hobbs, N., Kohli, P., Kleywegt, G., Birney, E., Hassabis, D., Velankar, S.: AlphaFold Protein Structure Database: massively expanding the structural coverage of protein-sequence space with high-accuracy models. Nucleic Acids Research 50(D1), 439–444 (2021) 10.1093/nar/gkab1061

[28] Zhang, Y., Skolnick, J.: Scoring function for automated assessment of protein structure template quality. Proteins: Structure, Function, and Bioinformatics 57(4), 702–710 (2004)

[29] Sen, Z.: Innovative Trend Methodologies in Science and Engineering. Springer, ??? (2017)

[30] Pedregosa, F., Varoquaux, G., Gramfort, A., Michel, V., Thirion, B., Grisel, O., Blondel, M., Prettenhofer, P., Weiss, R., Dubourg, V., Vanderplas, J., Passos, A., Cournapeau, D., Brucher, M., Perrot, M., Duchesnay, E.: Scikit-learn: Machine learning in Python. Journal of Machine Learning Research 12, 2825–2830 (2011)

[31] Hutchinson, E.G., Thornton, J.M.: A revised set of potentials for beta-turn formation in proteins. Protein Science 3, 2207–2216 (1994)

[32] Reeder, P.J., Huang, J.S. Y. M. and Dordick Bystroff, C.: A rewired green fluorescent protein: folding and function in a nonsequential, noncircular gfp permutant. Biochemistry 49, 10773–10779 (2010)

[33] Maciejewski, M.W., Schuyler, A.D., Gryk, M.R., Moraru, I.I., Romero, P.R., Ulrich, E.L., Eghbalnia, H.R., Livny, M., Delaglio, F., Hoch, J.C.: Nmrbox: a resource for biomolecular nmr computation. Biophysical journal 112(8), 1529–1534 (2017)

[34] Vosegaard, T.: Fast simulations of multidimensional nmr spectra of proteins and peptides. Magnetic Resonance in Chemistry 56(6), 438–448 (2018) 10.1002/mrc.4663 https://analyticalsciencejournals.onlinelibrary.wiley.com/doi/pdf/10.1002/mrc.4663

