## Supplemental Information for "NMR Spectroscopy for the Validation of AlphaFold2 Structures"

#### 1 Additional Training Details

In this section, we provide further details on training our support vector machines. Calculating CS and DS require values for the parameters $\epsilon_{CS}$ and $\epsilon_D$. $\epsilon_{CS}$ and $\epsilon_D$ are C$\alpha$ distances that provide cutoffs in the calculation of CS and DS. They were determined by a small grid search over a realistic range of parameters. For the distance score cutoff value, $\epsilon_D$, we calculated the correlation between DS and the TM-score over the training data for various values of $\epsilon_D$, see Figure 1. We expect this correlation between DS and TM-score to be negative, so we selected the value of $\epsilon_D$ that gives the greatest negative correlation. As seen in Figure 1, the best performing value was $\epsilon_D = 10$ Å.

Similarly, for the contact score threshold $\epsilon_{CS}$, we performed a grid search over the values 6, 9, 12, and 15 Å, and measured the correlation between the contact heuristic and the TM-score to find that 12 Å was optimal, Figure 2. These grid searches were performed before the full dataset had been collected and without the mismatched data. We chose to use these values for later calculations using both the original and augmented datasets to prevent over fitting of the parameters.

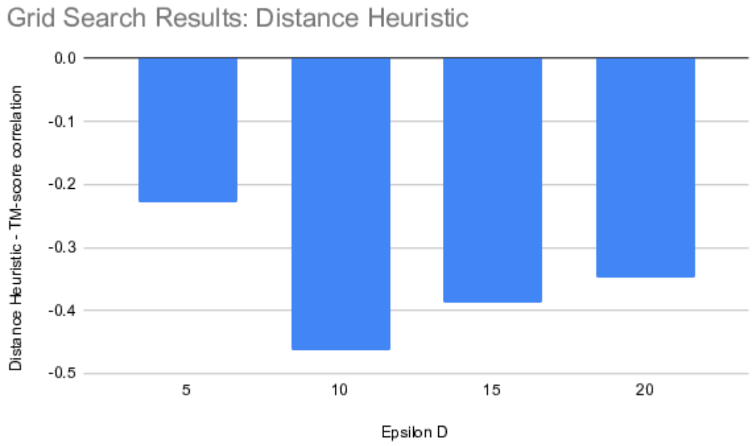


**Fig. 1**: Grid search over the distance score cutoff values shows that $\epsilon_D = 10$ Angstroms provides the greatest negative correlation between the distance heuristic and TM-score.

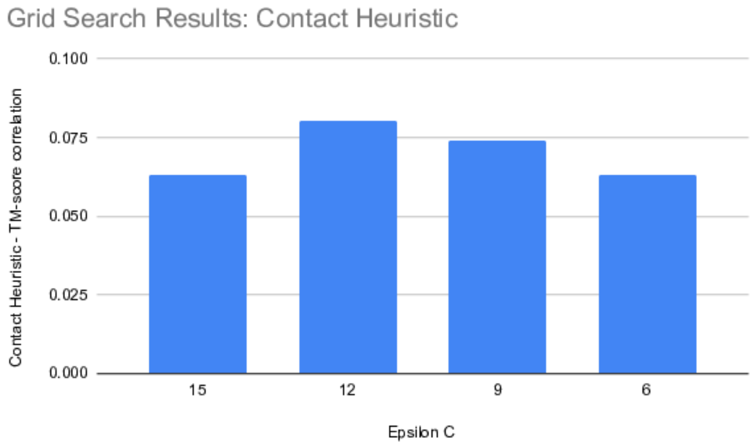


**Fig. 2**: Grid search over contact score cutoff values shows that $\epsilon_{CS} = 12$ Å provides the greatest correlation between the contact heuristic and TM-score.

#### 2 Comparison of Simulated versus Experimental NOESY Constraints

In our compiled dataset, only 117 members had experimental NOESY data thus requiring us to simulate NOESY data based upon NMR determined structures and the assigned chemical shifts. Simulated data were also produced for the dataset members with experimental data allowing us to test how well we recapitulated the experimental data. Histograms of these comparisons are below. We find that 91 $\pm$ 11% of the simulated peaks match those seen in the experimental data, while 9 $\pm$ 11% are new to the simulated data. As mentioned in the description of the simulation algorithm, some of these peaks could be due to unrealistic distances, thus providing artifactual peaks to our lists. Finally, 36 $\pm$ 11% of the peaks seen in the experimental data are missing from the simulated. This demonstrates that our simulation protocol is very conservative and in no way produces idealized data.

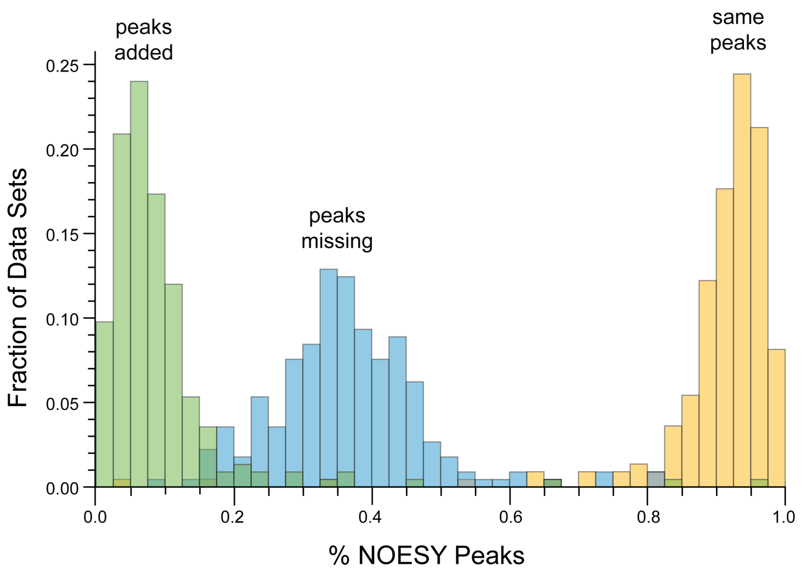


**Fig. 3**: Yellow: Histogram of percent of NOESY peaks that match between the experimental and simulated data. Blue: Histogram of the percent of peaks missing from the simulated data set compared to the experimental. Green: Histogram of the percent of peaks in the simulated data that are not seen in the experimental.

#### 3 Recall, Precision, and F-measure

Other methods besides our heuristics have been used to compare NMR data with proposed structural models. In particular, Huang et al. use recall, precision, and F-measure (RPF) to compare potential NOESY peak assignments to structural models.[1] RPF generates sets of proton pairs from both an input structure and an input NOESY spectrum, then applies the notions of recall, precision, and F-measure from information retrieval theory to compare the accuracy and completeness of the proposed NOESY assignment to the proposed structure. Specifically, recall is the percent of true HH pairs identified by the spectrum, precision is the distance-weighted percent of identified HH pairs that are true, and F-measure divides the product of the two by the sum.

We use RFP to analyze our dataset, but note that is different from that of Huang et al. in that we compare residue contacts and not HH pairs. We calculate our RPF values as follows:

1. Calculate a binary Potential Contact Map ($M_b$), by finding all pairs of residues $i$ and $j$ which have at least one potential contacts between them, and setting entry $M_{i,j} = 1$. All other entries are 0.
2. Calculate the AlphaFold Distance Map ($D$) by taking the distance between the $C_\alpha$s in each residue. Additionally, we calculate a binary AlphaFold Contact Map ($D_b$) by setting residues to be in contact when they are within $\epsilon_{CS}$, the threshold defined in Section **??**.
3. According to the RPF paper, we calculate the true positive rate ($TP$) as the sum over the element-wise product between $M_b$ and $D_b$. We also calculate a weighted true positive rate ($TP_w$) as the sum over the element-wise product between $M_b$ and $1/D^6$.
4. Then:
   - Recall $= TM/sum(M_b)$
   - Precision $= TM_w/sum(1/D^6)$
   - F-measure = 2*Recall*Precision/(Recall + Precision)

The correlation of these RFP values with our heuristics, PAE, and TM-score are calculated as before. Figure **??** shows the precision and F-measure as a function of TM-score over the augmented dataset. As before, the original data are blue, while the mismatched data are red. Figure 4) shows the values of the correlation and train additional SVMs (shown in Table 1.

| | Recall | Precision | F-measure | PAE | CS | DS | TM-score |
|---|---|---|---|---|---|---|---|
| **Recall** | 1.00 | 0.54 | 0.99 | -0.16 | 0.66 | 0.03 | -0.23 |
| **Precision** | 0.51 | 1.00 | 0.62 | 0.16 | 0.32 | 0.38 | -0.20 |
| **F-measure** | 0.99 | 0.59 | 1.00 | -0.13 | 0.66 | 0.07 | -0.24 |
| **PAE** | N/A | N/A | N/A | 1.00 | -0.30 | 0.26 | -0.46 |
| **CS** | 0.66 | 0.21 | 0.64 | N/A | 1.00 | -0.26 | 0.03 |
| **DS** | -0.29 | 0.08 | -0.26 | N/A | -0.68 | 1.00 | -0.28 |
| **TM-score** | 0.19 | 0.05 | 0.18 | N/A | 0.49 | -0.73 | 1.00 |

**Fig. 4**: Spearman's rank correlations between heuristics and TM-score, with recall, precision and F-measure included. Large positive values are in red and large negative values are in blue. Correlations presented in the upper triangle are from the original dataset, while the lower triangle comes from the dataset augmented with mismatched experiments.

Figure 4 shows that Recall, Precision, and F-measure are highly correlated with each other as one would expect from their definition. The CS is strongly correlated with Recall and F-measure and more weakly correlated with Precision. DS is primarily correlated with Precision and negatively correlated with CS, as was shown in the example cases above. While Recall and F-measure essentially measure the same characteristics of our dataset as indicated by a correlation coefficient of 0.99, the other heuristics manage to capture other relevant aspects of the data. Comparing the correlations above and below the diagonal, we see that the RPF scores perform as expected when identifying very good versus very bad structures but struggle when needing to distinguish structures that have fold similarity with the true structure but are not accurate.

Using the RPF metrics as an additional grouping of inputs for our SVM training, we get four new SVMs. These are compared with the original results in Table 1, showing that the RPF metrics are not as good on their own, but do benefit the combined SVM. Using all inputs together gives the best results, resulting in a balanced accuracy of 84%, with consistent structures identified at a rate of 89% and inconsistent structures at a rate of 79% in the test dataset.

### 4 LoTOP Purification

LoTOP was purified as described in the Materials and Methods. Size exclusion chromatagraphy shows single monomeric species. SDS page gel shows the purity of the NMR samples.

| Inputs | Accuracy (%) | | | |
|---|---|---|---|---|
| | Neg ID | Pos ID | Total | Balanced |
| RPF only [1] | 61.43 | 83.33 | 81.10 | 72.38 |
| PAE only | 40.00 | **86.73** | **81.98** | 63.37 |
| CS-DS only | **67.14** | 81.88 | 80.38 | **74.51** |
| RPF-PAE | 65.71 | 87.86 | 85.61 | 76.79 |
| CS-DS-RPF | 71.43 | 88.35 | 86.63 | 79.89 |
| CS-DS-PAE | 75.71 | 81.07 | 80.52 | 78.39 |
| All Inputs | **78.57** | **89.00** | **87.94** | **83.78** |
| Augmented Data | 92.07 | 90.23 | 91.21 | 91.15 |

**Table 1**: Comparison of accuracy of SVMs with different inputs on the binary classification task of identifying "consistent" (TM-score $> 0.5$) and "inconsistent" (TM-score $< 0.5$) structures using the original data set; *i.e.*, those without the mismatched data. Negative and Positive ID rates refer to the percent of "inconsistent" or "consistent" structures correctly identified. The SVMs were all trained with balanced class weights, therefore we include the balanced accuracy to demonstrate how well each model has achieved the goal of equally identifying positive and negative samples. The types of inputs used are labeled in the first column. The bottom line shows the results from training on the augmented data set in which the SVM was given only the CS and DS as inputs.

### 5 Design of LoTOP

LoTOP was designed by permuting the secondary structure elements of TOP7 thus simplifying its topology. This is shown in the figure.

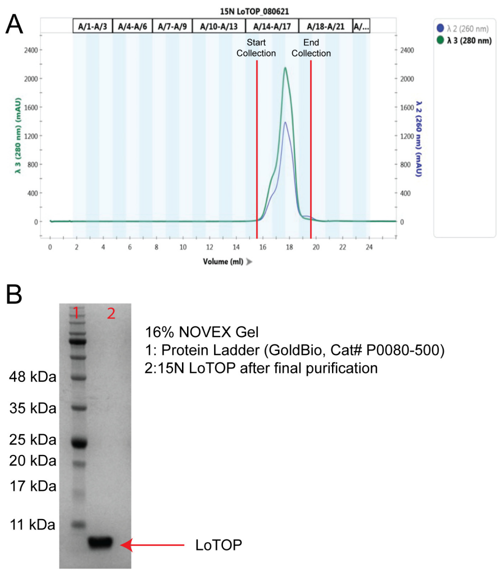


**Fig. 5**: Purification of LoTOP. A. Size Exclusion Chromatogram of $^{15}$N labeled LoTOP showing it to be a single monomeric protein. B. SDS page gel of $^{15}$N labeled LoTOP.

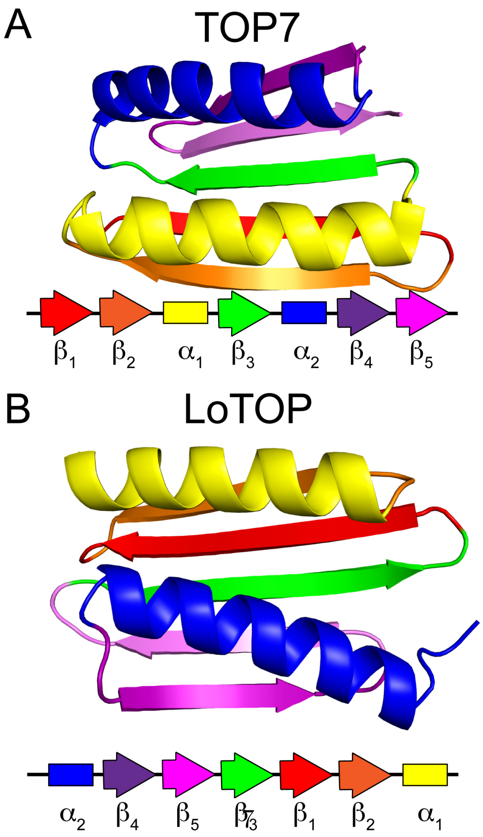


**Fig. 6**: Top7 and LoTOP structure and topology. A. The structure and topology of TOP7. B. The designed structure and topology of LoTOP.

### 6 LoTOP Stability

The stabiltiy of LoTOP was determined by measuring the change in the CD spectrum at 227 nm as the denaturant guanidinium hydrochloride was tritrated. The ΔG of unfolding was found to be 12.1±0.4 kcal/mol.

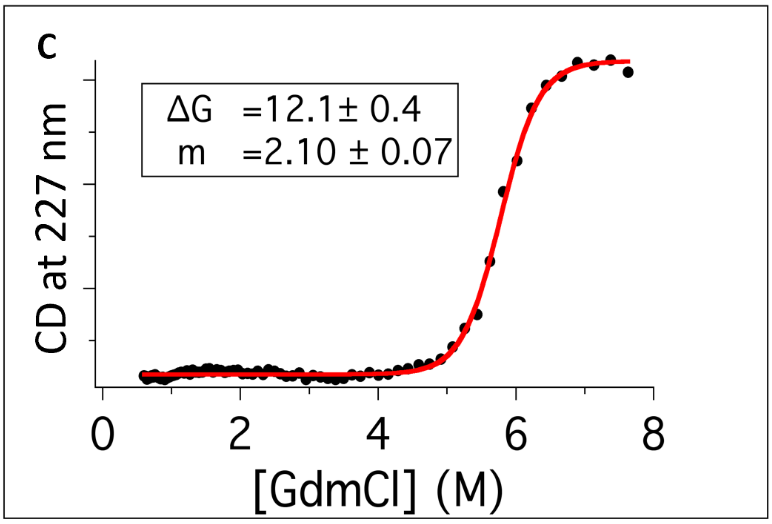


**Fig. 7**: Guanidinium hydrochloride titration of LoTOP.

#### 7 LoTOP NMR

$^{15}$N HSQC of LoTOP with HN assignments. Assignments were performed with standard triple resonance techniques.

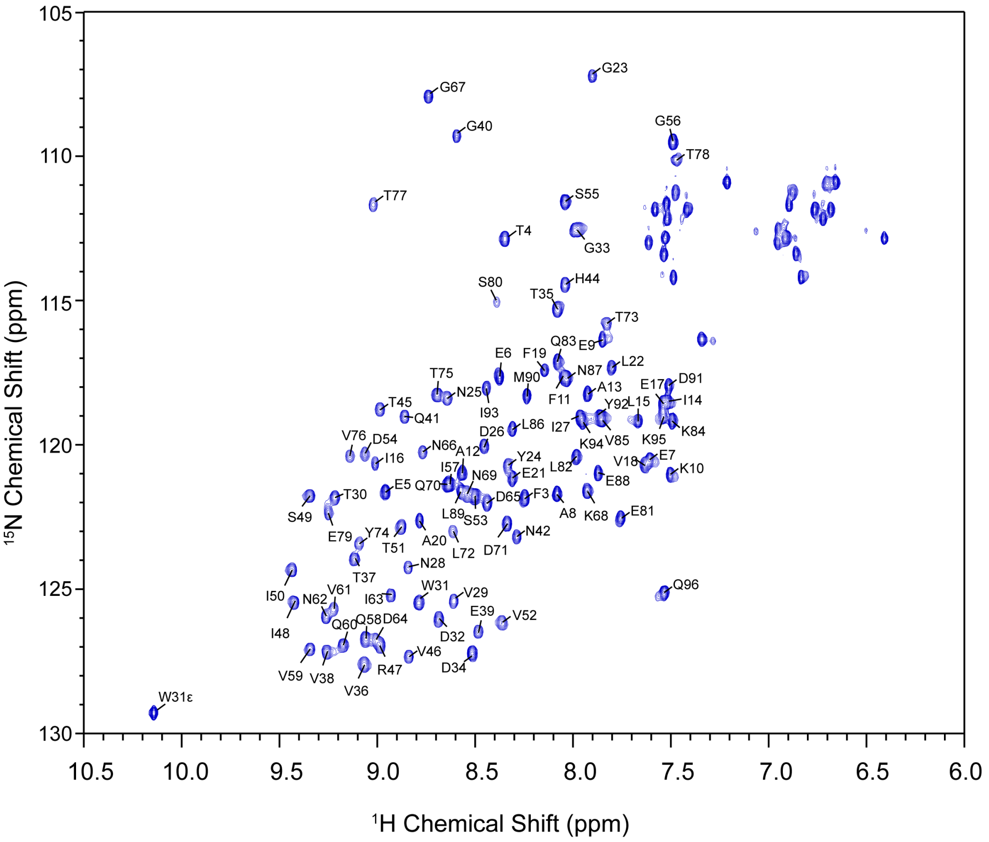


**Fig. 8**: $^{15}$N HSQC of LoTOP with HN assignments

#### References


[1] Huang, Y.J., Powers, R., Montelione, G.T.: Protein nmr recall, precision, and f-measure scores (rpf scores): Structure quality assessment measures based on information retrieval statistics. Journal of the American Chemical Society **127**(6), 1665–1674 (2005) https://doi.org/10.1021/ja047109h https://doi.org/10.1021/ja047109h. PMID: 15701001
